# Motivational context determines the impact of aversive outcomes on mental effort allocation

**DOI:** 10.1101/2023.10.27.564461

**Authors:** Mahalia Prater Fahey, Debbie M. Yee, Xiamin Leng, Maisy Tarlow, Amitai Shenhav

## Abstract

It is well known that people will exert effort on a task if sufficiently motivated, but how they distribute these efforts across different strategies (e.g., efficiency vs. caution) remains uncertain. Past work has shown that people invest effort differently for potential positive outcomes (rewards) versus potential negative outcomes (penalties). However, this research failed to account for differences in the context in which negative outcomes motivate someone - either as punishment or reinforcement. It is therefore unclear whether effort profiles differ as a function of outcome valence, motivational context, or both. Using computational modeling and our novel Multi-Incentive Control Task, we show that the influence of aversive outcomes on one’s effort profile is entirely determined by their motivational context. Participants (N:91) favored increased caution in response to larger penalties for incorrect responses, and favored increased efficiency in response to larger reinforcement for correct responses, whether positively or negatively incentivized.

**Statement of Relevance:** People have to constantly decide how to allocate their mental effort, and in doing so can be motivated by both the positive outcomes that effort accrues and the negative outcomes that effort avoids. For example, someone might persist on a project for work in the hopes of being promoted or to avoid being reprimanded or even fired. Understanding how people weigh these different types of incentives is critical for understanding variability in human achievement as well as sources of motivational impairments (e.g., in major depression). We show that people not only consider both potential positive and negative outcomes when allocating mental effort, but that the profile of effort they engage under negative incentives differs depending on whether that outcome is contingent on sustaining good performance (negative reinforcement) or avoiding bad performance (punishment). Clarifying the motivational factors that determine effort exertion is an important step for understanding motivational impairments in psychopathology.

## Introduction

Whether writing a paper or studying for a final exam, achieving our goals depends critically on our ability to generate sufficient motivation to carry out the tasks before us. When considering what leads people to fail to achieve a given goal, researchers and laypeople alike often focus on whether the promised rewards were sufficient (or sufficiently salient) to overcome the cost of engaging in the required effort (Emanuel et al., 2022; Fishbach et al., 2010; Gollwitzer & Sheeran, 2006). However, it has long been understood that motivation can arise as much or more from the potential negative outcomes of *failing* to engage effort (e.g. reprimands, loss of earnings) as it does the positive outcomes of *succeeding* to engage effort (e.g. praise, increase of earnings) (Atkinson, 1957; Yee et al., 2022; Yee & Braver, 2018). Recent work has sought to formalize the process by which positive and negative incentives are integrated to determine effort allocation (Leng et al., 2021; Ritz et al., 2022; Yee et al., 2022), and has predicted conditions under which these outcomes should contribute not only to different amounts of effort but also different *kinds* of effort. However, while this and other research have begun to disentangle how negative incentives differ from positive incentives in determining mental effort motivation, it has yet to address how negative incentives differ from *one another* depending on the context in which they are used to motivate effort.

Models of decision-making provide a useful framework for disentangling the ways in which expected outcomes determine effort investment into a task (Shenhav et al., 2017). For instance, to maximize our rate of expected reward, we may choose to adjust our cognitive control strategy to achieve greater speed and/or greater accuracy (Bogacz et al., 2006). We recently showed that people favor different control strategies depending on the type of incentive being varied, specifically whether it is the positive consequences for good performance or the negative consequences for poor performance (Leng et al., 2021). By manipulating the amount of monetary reward for correct responses and the amount of monetary losses incurred for incorrect responses, we found a dissociation whereby larger rewards motivated participants to respond more efficiently (faster with little sacrifice to accuracy, resulting in an overall increase in productivity on the task) and larger penalties motivated them to respond more slowly but also more accurately (resulting in an overall decrease in productivity). We showed that these dissociable patterns of behavior matched the predictions of a reward-rate maximizing model of control allocation, under the key assumption that participants are able to distribute their effort across multiple different types of control (in this case, leading to differential adjustments of evidence accumulation rate and decision threshold) (cf. Ritz et al., 2022).

This work demonstrated that rewards and penalties promote distinct control strategies, but in doing so it conflated two differences between those incentives. In addition to differing in valence (positive vs. negative), these incentives also differed in their *motivational context*, that is, whether they served as *reinforcement* (of correct responses) or *punishment* (of incorrect responses) (Yee et al., 2022). It is therefore unclear to what extent these dissociable control strategies can be attributed to differences in incentive valence, incentive type (reinforcement vs. punishment), or both. Addressing this question requires comparing conditions in which an equally valenced outcome (e.g., aversive) serves as either a punishment (discouraging poor performance, as in the previous study) or as reinforcement (encouraging good performance, as remains to be tested). This comparison might reveal that the type of control strategy favored is primarily determined by incentive valence, for instance with negative reinforcement promoting accuracy over speed (as previously observed when varying punishment). Alternatively, it may reveal that the favored control strategy is primarily determined by motivational context, for instance with negative reinforcement largely promoting more efficient responding (as previously observed when varying reward, i.e., positive reinforcement). In addition to disambiguating our previous findings, the outcome of such a comparison could shed important light on past research on motivation and cognitive control, where both the valence and type of incentive have varied and resulting behaviors have been heterogenous (Braem et al., 2013; Cubillo et al., 2019; Levy & Schiller, 2021; Ličen et al., 2016; Mobbs et al., 2020; Yee et al., 2015, 2021).

To formally test the extent to which control strategies reflect *incentive valence* or *motivational context*, we designed a novel variant of the Multi-Incentive Control Task. In this new version of the task, monetary loss could occur as a consequence of performing poorly (penalties incurred for each incorrect response, as before) or as a consequence of failing to perform well (losses that could be avoided with each correct response). We varied the magnitude of penalty (punishment) and loss-avoidance (negative reinforcement) orthogonally to one another, and separately from potential rewards for each correct response (positive reinforcement). We found that participants adjusted their control strategy based on the magnitude and the motivational context of a given incentive. Specifically, aversive outcomes led participants to either increase their caution or their speed/efficiency depending on whether the outcomes were varied in the context of a punishment or a (negative) reinforcement, respectively; changes in task performance in the face of increasing negative reinforcement mirrored those observed with increasing positive reinforcement. These dissociable strategies were well-accounted for by a model that configures control by maximizing reward rate and minimizing effort costs. Collectively, this work sheds new insight into the process by which people integrate diverse inputs from their environment to determine how much and what kind of mental effort to exert.

## Methods

### Participants

The study was approved by Brown University’s Institutional Review Board, and participants provided informed written consent and were compensated in cash for their participation. We recruited 117 participants on Prolific, who indicated they were within the United States, had normal or corrected to normal vision, no dyslexia or color blindness, and were currently attending university. This dataset was collected as part of a broader study goal where active attendance at university was a pertinent requirement. We excluded 20 subjects for not understanding the task correctly (as determined across 12 quizzes before starting and during the task). We excluded 1 participant for displaying button-smashing behavior (quantified as responding faster than 250ms on more than 30% of responses). We additionally excluded 3 participants who did not complete the study (did not reach the end of the task) and participants who did not provide enough behavioral responses for data analysis (responded to less than 80% of intervals). Our final sample consisted of 91 participants (43 females and 48 males; mean age 23.35; 12.10% Asian, 8% Black/African American, 58.24% Caucasian, 9% Hispanic/Latinx, 13.2% Mixed). Participants were paid $8.00 per anticipated hour on the task and earned on average $4.42 (sd: 0.84) in bonus money. We conducted a power analysis (Murayama et al., 2022) on a separate data set in which we had run participants on the Collector Game (See Multi-Incentive Control Task) to determine the number of subjects to include to find an interaction with reinforcement magnitude and penalty magnitude at the level of the correct responses per second during the interval and reaction time at the level of trial. The power analysis at the level of the interval suggested that with 100 subjects we would be powered with 0.87 to find the effect with an effect of 0.31. The power analysis on reaction time suggested we would be powered with 0.95 to find the effect with an effect of 0.36.

### Multi-Incentive Control Task

To test whether the motivational context of negative outcomes guides dissociable strategies for cognitive control allocation, we adapted and modified the Multi-Incentive Control Task developed in Leng et al. (2021), and included a cover story or whether participants could earn monetary rewards in the form of gems or lose monetary rewards in the form of bombs. The task consists of “turns” during which they had intervals of Stroop trials varying between 6-9 seconds, which ensured that participants would not expect a fixed number of stimuli per interval. During the interval, participants could complete as many cognitively demanding Stroop trials as they wished within a fixed time interval. Participants were asked to respond to the ink color (red, yellow, green, blue) of a color word (‘RED’, ‘YELLOW’, ‘GREEN’, ‘BLUE’) by pressing the corresponding key on their keyboard. Trials consisted of congruent stimuli where the ink color and the color word were in agreement (e.g. GREEN in green ink) and incongruent stimuli where the ink color and the color word were not in agreement (e.g. GREEN in blue ink). An important feature of the Stroop task is it requires cognitive control to accurately respond to the ink color while inhibiting automatic reading of the color word, and therefore successful performance of the Stroop task would reflect increased attention to the changing stimulus (MacLeod, 1991; Stroop, 1935). The overall percent of congruent to incongruent stimuli was 45.80%, which could vary within an interval due to the self-paced display of stimuli.

Participants completed two “games” of the Stroop task (counterbalanced) that varied in the motivational context of expected outcomes for correct and incorrect task performance (illustrated in Figure 1a). At the start of the experiment, participants were initially endowed with $12.00 of bonus money which was converted into 1200 gems for the game and added to a bank. In the Collector Game, participants were instructed that correct responses would increase this endowment (positive reinforcement). Conversely, in the Protector Game, correct responses would reduce a large potential loss (negative reinforcement). In both games, incorrect responses would incur a penalty of a monetary loss that would reduce this endowment (punishment). Critically, this framing of whether participants are instructed to pursue the task goal of maximizing monetary bonus versus minimizing monetary loss provides an important distinction that can allow us to dissociate and quantify how negative outcomes can drive correct and incorrect performance on a cognitive control task (and whether they would bias dissociable strategies for mental effort allocation).

**Figure 1.**
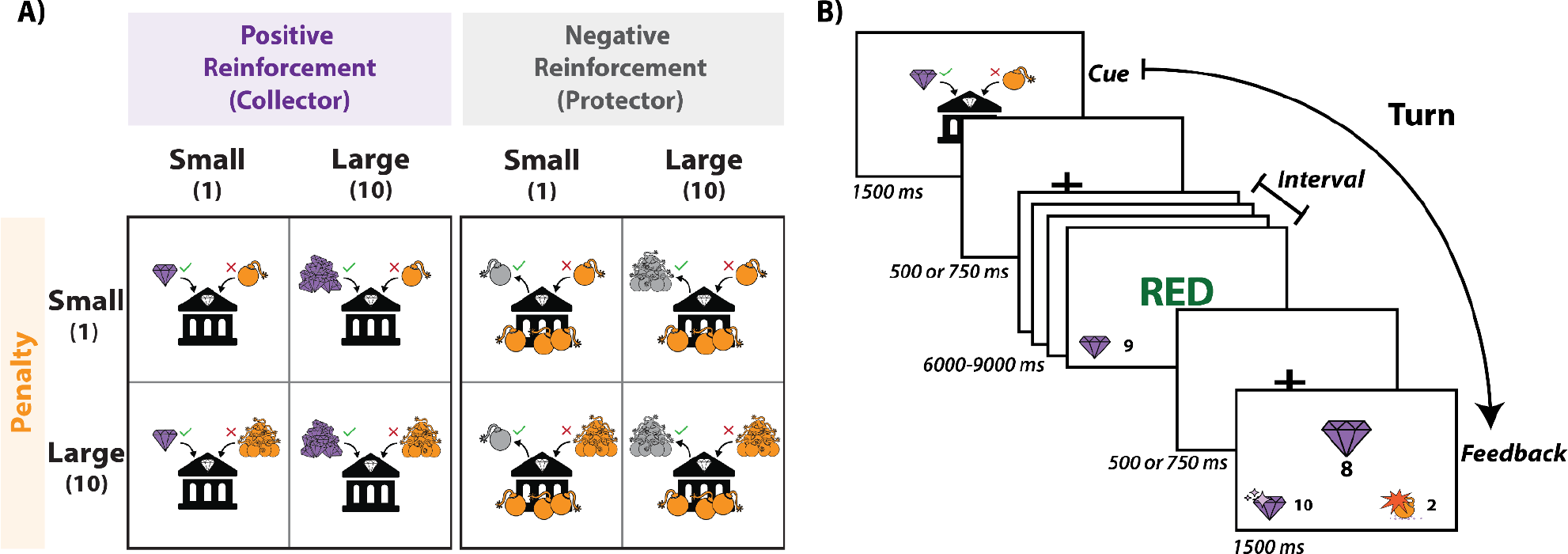
Multi-Incentive Cognitive Control Task. A) *Incentive Cues*. The magnitude of reinforcement for correct responses was either small (1) or large (10). In the Collector Game, participants could add either 1 or 10 gems to their bank with each correct response. In the Protector Game (negative reinforcement), participants were threatened with a potential loss of 300 gems and could remove either 1 or 10 bombs with each correct response, which would reduce the amount of gems they would lose from their bank. Across both games, the punishment for each incorrect response was either small (1 bomb) or large (10 bombs). B) *Example Turn*. Each turn began with a cue (1.5 s) indicating the magnitude of the reinforcement for each correct response and the penalty for each incorrect response. This was followed by a variable interstimulus interval (.5 or .75 s) followed by the interval start. During the interval (6-9 s), participants could respond as fast or slow as they wanted to the Stroop stimuli. After each response, a brief fixation cross would appear (.25 s) followed by the next Stroop stimulus until the interval ended. A tracker displayed either the number of gems added or bombs removed for each correct response along with the number of bombs added for each incorrect response. At the end of the interval, participants saw the net earnings based on their correct and incorrect responses. In the Collector Game (positive reinforcement), this was equivalent to the number of gems earned for correct responses subtracting the number of bombs detonated for incorrect responses. In the Protector Game (negative reinforcement), this was equivalent to the remaining bombs from the initial 300 after subtracting the number of bombs removed for correct responses and adding the number of bombs detonated for incorrect responses.

At the start of each turn, participants viewed a visual cue (1.5 s) which indicated the magnitude of the reinforcement and penalty for the upcoming interval. We varied the magnitude of reinforcement for correct responses (1 vs. 10) and the magnitude of the penalty for incorrect responses (1 vs. 10) within each game. Each game contained 4 blocks, resulting in 8 blocks total (Figure 1). The games were split into 4 task blocks of 15 turns (block order was randomized across participants). Within each block, the magnitude of one dimension of the cue was held constant while the magnitude of the other dimension was varied. For example, in a high reinforcement block, all turns within that block would have a large reinforcement (10), but turns within that block randomly varied between small (1) or large (10) penalty. During the interval, participants could respond to as many or as few Stroop stimuli as they liked. After each response, a brief fixation cross appeared (.25 s), and then the next stimulus would appear. During the interval, a tracker separately displayed the cumulative reinforcement (positive: gems added, negative: bombs removed) and punishment (both: bombs added) incurred. At the end of the turn, participants received feedback (1.5 s) during which the tracker remained on the screen along with the net summation of the total reinforcement and punishments incurred during the interval. Participants were informed that within each block one of the turns would be selected to be part of their final bonus (4 per game and 8 in total). The experiment was implemented using the Psiturk framework.

### Procedure

After providing consent and completing surveys participants began the task. Participants completed several rounds of practice to become familiar with the task. First, they learned the keymapping between the colors (‘D’-red, ‘F’-yellow, ‘J’-green, ‘K’-blue) by completing 80 trials in which they had to respond to a colored string of “XXXXXX” within 2 seconds. After each trial, they received feedback as to whether they were correct, incorrect, or too slow. Next, participants were quizzed on comprehension of Stroop instructions before completing 60 trials of Stroop practice which was also timed, and received the same feedback after each response. Participants then learned about the self-paced intervals and completed 4 practice turns. After learning the basic task structure, participants were then introduced to the first of the two games (game order was counterbalanced), through which they learned the magnitude of gain or loss avoided for correct responses and the magnitude of punishment for incorrect responses. Participants then completed two quizzes. The first assessed their comprehension of the cues for each incentive condition. The second presented hypothetical correct and incorrect responses made during a turn in each of the 4 incentive conditions and asked them to indicate what the resulting outcome would be at the end of the turn. Upon successfully completing the quizzes participants completed two practice turns. After the practice, participants completed the 4 blocks of the first game. Before each block participants were shown the incentive cues and were quizzed for comprehension of the cue and what the resulting outcome would be at the end of turns given that incentive cue. After finishing the first game, participants were introduced to the incentive conditions of the second game, completed the two quizzes and practice turns, followed by the 4 blocks of the second game. Again, before each block participants were quizzed on what the cues meant and what the outcome would be at the end of the turn given hypothetical responses. After participants completed both games, participants completed a questionnaire in which they viewed the incentive cues for each game and reported Likert ratings on a scale from -5 to 5 across 6 different dimensions (effort, motivation, pleasantness, attention, arousal, difficulty).

### Collector Game (Positive Reinforcement and Punishment)

During the Collector Game, participants could collect additional gems to add to their initial endowment. Each correct response added gems to the participants’ banks and each incorrect response added bombs to their banks. At the start of each turn, the cue indicated whether the participant would earn a small or large number of gems for each correct response, as well as whether they would add a small or large number of bombs for each incorrect response. During the interval, the tracker displayed the cumulative number of gems they added to their bank for each correct response as well as the bombs they added for each incorrect response. At the end of each turn, participants received feedback on the net gems they earned or lost (gems added - bombs added). At the end of the game, 1 of the 15 turns in each block was randomly selected to determine the final bonus to be added to the endowment.

### Protector Game (Negative Reinforcement and Punishment)

This Protector Game was the same as the Collector Game, except that participants protected the gems in their initial endowment, which were threatened by 300 bombs that were added to their bank at the start of each turn. Each correct response removed bombs from the participant’s bank and each incorrect response added additional bombs to their bank. At the start of each turn, a cue indicated whether the participant would remove a small or large number of bombs for each correct response, as well as whether they would add a small or large number of bombs for each incorrect response. During the interval, the tracker displayed the cumulative number of bombs they removed from the initial 300 bombs that were added at the start of the turn for each correct response as well as the bombs they added for each incorrect response. Participants received feedback at the end of the turn on the net number of bombs remaining in the bank (Initial 300 bombs - bombs removed + bombs added). At the end of the game, 1 of the 15 turns in each block was randomly selected to determine the final bonus to be added to the endowment.

### Behavioral Analyses

An innovative feature of the self-paced design is that it allows us to analyze performance at the level of the interval as well as at the individual trials. We analyzed participants’ performance at the level of the turn by fitting a linear mixed model (lme4 package in R; Bates et al., 2015) to estimate the correct responses per second as a function of the contrast coded reinforcement magnitude (large reinforcement = 1, small reinforcement = -1), penalty magnitude (large punishment = 1, small punishment = -1) This model concatenates the data from both games allowing us to include reinforcement valence (positive = 1, negative = -1) and the full 3-way interaction between reinforcement magnitude, penalty magnitude, and reinforcement valence. The model also controlled for average congruency of the interval (z-scored; Congruent = 1, Incongruent = 0), interval length (z-scored), interval number the entire session (z-scored), game order (dummy coded; loss-then-gain = 1, gain-then-loss = 0), the interaction between game order and reinforcement valence, sex (female = -1, male =1), and age (z-scored). All of these variables are contrast or dummy coded in the same manner in all subsequent mixed models, and asterisks indicate interactions between variables. To determine the random effects for the current and all subsequent models, we first fit the maximally specified random effects structure interacting valence, penalty, and reinforcement. If the maximally specified model failed to converge, we then reduced the random effects structure by removing the random effect that explained the least amount of variance until we reached convergence (Barr et al., 2013). For the current model, our final random effects structure included reinforcement magnitude, penalty magnitude, and reinforcement valence.

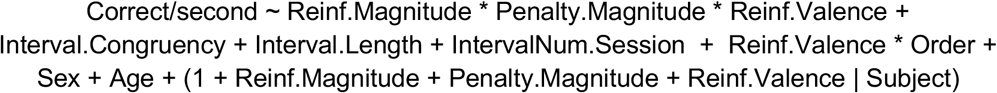

#### Correct Responses Per Second

When we examined correct responses per second within each game separately, we used the same fixed effects structure, except without the reinforcement valence. The models controlled for interval congruency (z-scored), interval length (z-scored), interval number within the specific game (z-scored), game order, sex, and age. Our final random effects structure in the Collector Game included the main effects for reinforcement magnitude and penalty magnitude. In the Protector Game, the model converged with the maximal random effects structure interacting penalty magnitude with reinforcement magnitude but we reduced it to match the effect structure across games

Collector Game:

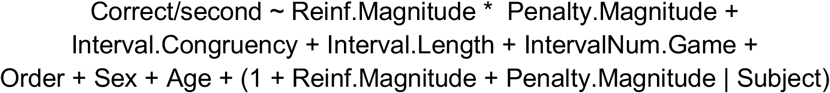

Protector Game:

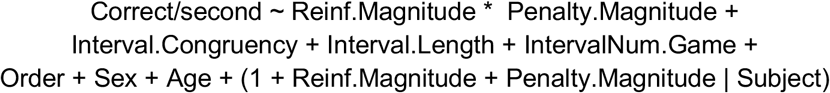

To examine how overall performance at the turn level was determined by adjustments in speed and accuracy, we fit linear mixed models to log-transformed reaction time (correct responses only) and generalized linear mixed effects models to accuracy, both at the trial-level.

#### Accuracy

In our accuracy model, we controlled for age, sex, congruency, interval length (z-scored), interval number in the entire session (z-scored), trial number in the interval (z-scored), as well as game order, and the interaction between game order and reinforcement valence. Our final random effects structure included reinforcement magnitude, penalty magnitude, and reinforcement valence.

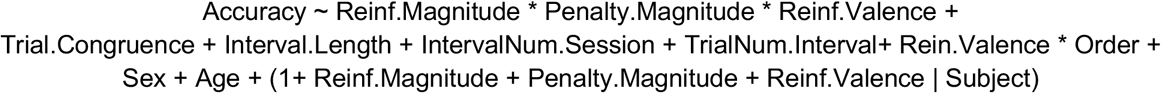

When we separately examined accuracy within each game, we used the same fixed effects structure, except without the reinforcement valence. We controlled for congruency, interval length (z-scored), interval number within the specific game (z-scored), trial number in interval (z-scored), game order, age, and sex. Our final random effects structure in the Collector and Protector Game included reinforcement magnitude and penalty magnitude.

Collector Game:

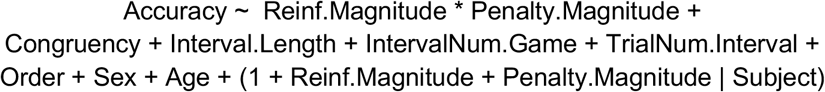

Protector Game:

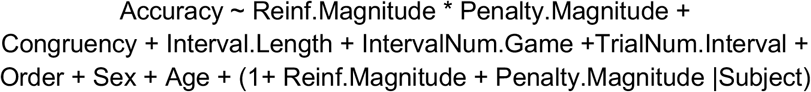

#### Reaction Times

In our reaction time model, we analyzed the log-transformed reaction time of accurate trials only. We controlled for trial congruency, interval length (z-scored), interval number in the entire session (z-scored), trial number in the interval (z-scored), game order, the interaction between game order and reinforcement valence, sex, and age (z-scored). Our final random effects structure included reinforcement magnitude, penalty magnitude, and reinforcement valence.

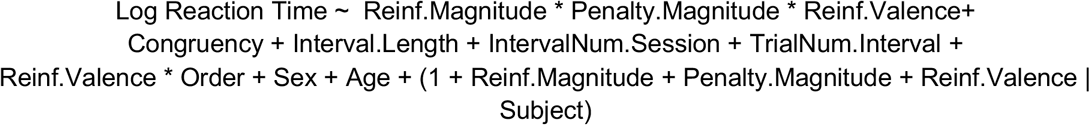

When separately examining log reaction time within each game, we used the same fixed effects structure, except without the reinforcement valence. We controlled for trial congruency, interval length (z-scored), interval number within the specific game (z-scored), trial number within the interval, game order, sex, and age (z-scored). Our final random effects structure in the Collector Game and Protector Game included reinforcement magnitude and penalty magnitude.

Collector Game:

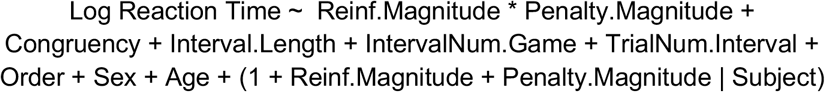

Protector Game:

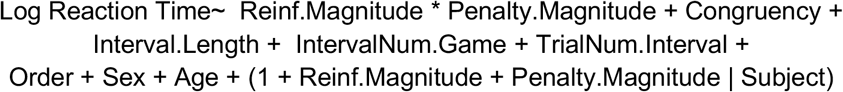

### Computational Modeling Analyses

#### Model Fitting

A Drift Diffusion Model (DDM) (Ratcliff & McKoon, 2008) was used to parameterize participants’ responses on the Stroop task as correct (e.g., responding to the ink color) or incorrect (e.g., responding to the word). Participants could either make an error by responding to the Stroop stimulus word (an automatic error) or by responding with one of the other 2 Stroop words (a random error). Given that we were interested in characterizing the decision in cognitive control allocation and to reduce noise in the model fitting process, we only included automatic errors (e.g., incorrectly responding to the word during an incongruent trial) and excluded random errors (e.g., incorrectly responding to a color not associated with the current stimulus). This did not significantly impact the dataset, since random errors only consisted of 4.26% of total trials. Next, we fit a regression model in the Hierarchical Drift Diffusion Model package (Wiecki et al., 2013) and performed Bayesian parameter estimation of drift rate and threshold parameters. As with our model-agnostic analyses, we fit models for each game separately as well as one that concatenated data across the two games. We estimated the parameters of drift rate and threshold as a function of reinforcement (large reinforcement = 1, small reinforcement = -1), and penalty (large punishment = 1, small punishment = -1). When fitting the models with data concatenated from both games together we included a regressor for the valence of the reinforcement (positive = 1, negative = -1). To account for congruency effects, the model for drift rate included trial congruency (congruent = 1, incongruent = 0). We ran 5 parallel chains for 12000 iterations each, discarded the first 8000 warm-up samples, and kept the remaining 4000 samples as posterior estimates (20000 samples total). The Gelman-Rubin convergence diagnostic Rhat (Gelman & Rubin, 1992) was used to assess model convergence (Rhats close to 1 indicates convergence; we considered a model successfully converged with Rhats <= 1.01 (Baribault & Collins, 2023).

Next, we performed model comparison to determine the best-fitting regression model. We ran 20 models and examined to what extent inclusion of intertrial variability, fixed starting point, or biasing for congruence improved our model fitting procedure (Table S4). Additionally, we accounted for the collapsing response deadline (i.e., decision threshold) associated with the reduced amount of time available to complete a given trial within each interval (Fengler et al., 2022; Palestro et al., 2018). For example, for a given 5-second interval, if the participant took 500 ms to respond to the first stimulus, when adding in the intertrial interval time of 250 ms, then the participant would have 4250 ms to complete additional trials within that interval. To approximate this interval-level collapsing bound, we included a variable called “scaled linear running time”, which provided the amount of time that had passed within each interval (z-scored). Nondecision time was fit as a free parameter. We performed model selection by examining whether including variables improved model fit based on the Bayesian predictive information criterion (BPIC) The best-fitting models included intertrial variability, a bias on the starting point determined by the congruency of the trial, and included running time on the threshold (BPICs are illustrated in Figure S2). We performed additional control analyses to rule out the possibility that the putative effect of valence (e.g., collector game/positive reinforcement vs. protector game/negative reinforcement) was confounded with game order (e.g., first vs. second game), and observed non-significant effects of game order when included in the model and with worse BPICs.

Posterior prediction checks were performed to validate the model fitting procedure (Wilson & Collins, 2019). We generated 500 independent samples from the posterior distribution of fitted parameters and then simulated the reaction time distribution for each posterior sample. The predicted reaction time distribution matches with the actual reaction time distribution and error rate for each condition (Figures S4, S6, S8).

### Reward Rate Model

To test our normative predictions of reinforcement and punishment effects on mental effort allocation, we used the reward rate model detailed in Equation 1 (Bogacz et al., 2006; Leng et al., 2021). We tested these predictions using the winning HDDM models that were fit to the data from each game separately. This model assumes the drift diffusion model as the underlying process by which participants respond to either the ink color (target) or the word (distractor). A key assumption of the model is that participants are able to maximize their overall reward rate within the interval by allocating their cognitive control, combining separate control strategies that tradeoff between increasing efficiency of processing visual stimuli (e.g., drift rate) and decision threshold in a multivariate manner (Ritz et al., 2022). The model assumes that participants adjust these combinations of drift rates and thresholds depending on the weight they place on the potential reinforcements for correct responses as well as the weight they place on potential punishments for incorrect responses. Specifically, our reward-rate model predicts that participants should increase their drift rate when they can earn greater reward or avoid a larger loss for a correct response, whereas they should increase their response threshold when they would receive a larger penalty for an incorrect response (Fig 3A).

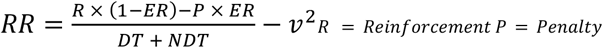

### Subjective Cue Ratings

We analyzed participants’ ratings of subjective experience of each cue by fitting a robust linear model (robustbase package in R, Koller & Stahel, 2011). We estimated participants’ ratings of their effort, motivation, pleasantness, attention, arousal, and difficulty as a function of the contrast coded reinforcement magnitude (large reinforcement =1, small reinforcement = -1), penalty magnitude (large punishment =1, small punishment =-1), and reinforcement valence (positive = 1, negative = -1), as well as their interactions. The models also controlled for age and sex. When we examined subjective experiences within each game separately, we used the same fixed effects structure, except without the reinforcement valence.

Next, we ran spearman correlations between all the cue ratings and observed high correlations between the motivation, effort, and attention ratings (Collector: *R*_*(362) Motivation-Effort*_ = 0.61, p<0.001; *R*_*(362) Motivation-Attention*_ =0.44, p<0.001; *R*_*(362) Attention-Effort*_ =0.67, p<0.001; Protector: *R*_*(362) Motivation-Effort*_ = 0.55, p<0.001; *R*_*(362) Motivation-Attention*_ =0.40, p<0.001; *R*_*(362) Attention-Effort*_ =0.64, p<0.001 Fig S9). Given their high correlations, we then created a composite motivation rating by taking the motivation, effort, and attention ratings of the cues.

## Results

To understand whether the context of aversive outcomes influences the selection of cognitive control strategies and to tease apart the distinct roles of *incentive valence* (positive versus negative) and *motivational context* (reinforcement versus punishment) in driving control allocation, participants play two games. For one of these games (the *Collector Game)*, participants could collect a reward for each correct response (symbolized by gems) and incur a penalty (monetary loss) for each incorrect response (symbolized by bombs). Across the experiment, we varied the size of the reward and penalty (1 vs. 10 gems/bombs for correct/incorrect responses). Participants were cued with the relevant performance contingencies prior to each task interval (Fig 1). The Collector Game was designed to match the the tasks used in Leng et al. (2021), capturing the impact of prospective positive outcomes in the form of reinforcement and the impact of negative outcomes in the form of punishment.

To test the hypothesis that aversive outcomes should engender distinct control strategies depending on whether they are being used to reinforce or punish behavior, we had participants perform another game (the *Protector Game*) that focused entirely on aversive outcomes, which varied across both motivational contexts. Participants started the game with an endowment (1200 gems) that came under threat on each turn. Specifically, each turn would start with the potential for losing up to 300 gems (symbolized with the equivalent number of bombs) and each correct response would serve to reduce that potential loss by destroying a certain number of bombs (Fig 1). Mirroring variability in positive reinforcement in the Collector Game, in the Protector Game each correct response could avoid either a small or a large loss (destroying either 1 or 10 bombs), entailing either a small or large level of negative reinforcement. As in the Collector Game, each incorrect response incurred a small or large penalty (1 vs. 10 bombs), and information about task incentives was cued at the start of a turn. We could then directly compare the performance promoted by the aversive penalties in both games and the positive reinforcement in the Collector game to the novel condition of negative reinforcement in the Protector game.

### Control adjustments dissociate based on motivational context rather than incentive valence

When varying the magnitude of both positive reinforcement and punishment (Collector Game), we replicated the behavioral dissociation previously observed (Leng et al., 2021): with larger rewards for correct responses, participants responded substantially faster (*F*_(1,86)_ = 84.55, p<0.001) and slightly less accurately (*Chisq*_(1)_= 4.25, p= 0.04) (Fig. 2A,B, Table S1), collectively resulting in them completing more correct trials per second (*F*_(1,88)_ = 62.78, p<0.001) (Fig S1, Table S1). With larger penalties for incorrect responses, participants responded more accurately (*Chisq*_(1)_= 40.96, p<0.001) but substantially slower (*F*_(1,83)_ = 49.12, p<0.001) (Fig. 2A,B, Table S1), collectively resulting in fewer correct responses per interval on average (*F*_(1,91)_ = 7.24, p=0.009) (Fig S1 /Table S1). Given that the conditions for punishment were identical across our two games, we expected that responses to penalties in one task would mirror the other. This was indeed the case: just as in the Collector Game reported above, when performing the Protector Game participants were slower (*F*_(1,83)_ = 35.91, p<0.001) and more accurate (*Chisq*_(1)_= 13.05, p<0.001) for larger penalties (Fig. 2A,B, Table S2). This again resulted in slightly lower rates of correct responding (*F*_(1,91)_= 5.39, p= 0.022) (Fig S1 /Table S2).

**Figure 2.**
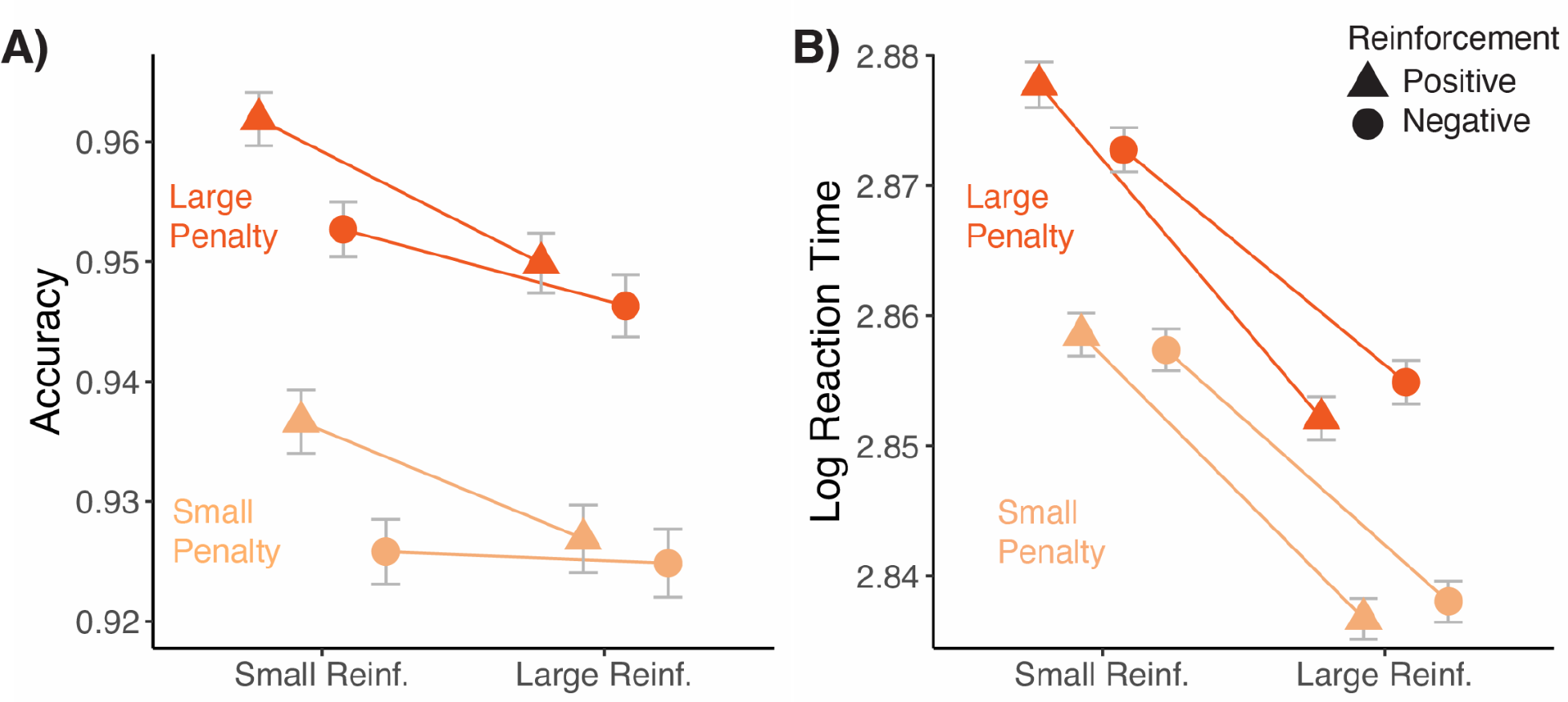
Effects of incentive magnitude, valence, and motivational context on performance. Effects of negative and positive reinforcement and punishment on overall task performance revealed that participants account for motivational context of aversive outcomes when determining cognitive control allocation. A) Participants were slightly less accurate for smaller positive reinforcements (left triangles) compared to larger positive reinforcements (right triangles). In both the Collector (triangles) and Protector (circle) games participants significantly increased their accuracy for higher penalties (dark orange) compared to smaller penalties (light orange). B) Participants were significantly faster for larger positive reinforcements (right triangles) compared to smaller positive reinforcements (left triangles). In both the Collector (triangles) and Protector (circle) games participants were significantly slower for higher penalties (dark orange) compared to smaller penalties (light orange). Crucially, when expecting higher negative reinforcement (circles), participants performed in a similar manner to positive reinforcement, and they responded faster and trended towards reducing their accuracy. Error bars reflect standard errors.

A crucial feature of the Protector Game is the inclusion of aversive outcomes across two motivational contexts (reinforcement vs. punishment). This provided us the unique opportunity to explicitly test whether behavioral responses to greater magnitudes of loss avoidance (with correct responding) would more closely mirror the effect of potential punishment (consistent with a general response to outcome valence) or the effect of potential rewards in the Collector Game (consistent with a motivational context-specific response). Our results are consistent with the second hypothesis. Participants were not slower and more accurate for greater loss avoidance, as would be expected if they responded to these aversive outcomes as they did punishment; instead, with greater magnitude of loss avoidance they responded much faster (*F*_(1,77)_= 34.05, p<0.001) and trended towards being less accurately (*Chisq*_(1)_= 3.34, p= 0.068, Fig. 2 A,B, Table S2), collectively resulting in a higher rate of correct responding (*F*_(1,91)_= 27.88, p< 0.001) (Fig S1 /Table S2). In other words, participants responded to greater loss avoidance in the Protector Game the same way they did to greater reward in the Collector Game, with the common factor being that both types of incentives *reinforced* correct responding.

To test whether participants responded differently to the two types of reinforcement (positive vs. negative), we analyzed task performance in the model that included concatenated data across both games in experiment. We found that participants were slower (F_(1,87)_= 63.88, p<0.001) and more accurate (*Chisq*_(1)_= 39.45, p<0.001) for larger penalties, whereas they were faster (*F*_(1,86)_= 84.45, p<0.001) and slightly less accurate (*Chisq*_(1)_= 4.57, p<0.033) for larger reinforcements (larger reward in the Collector game, larger loss avoidance in the Protector Game) (Table S3). Collectively, this resulted in higher rates of correct responding for larger reinforcements (*F*_(1,90)_= 79.74, p<0.001) and lower rates for larger punishments (*F*_(1,90)_= 10.17, p=0.002). When comparing the two types of reinforcement, we found that participants performing the Collector game were slightly slower (*F*_(1,87)_ = 2.92, p=0.091), but accuracy was similar (*Chisq*_(1)_= 0.241, p=0.623) (see also Table S3). We also found that reinforcement valence interacted with reinforcement magnitude in reaction time (*F*_(1,87)_ = 6.70, p=0.010) but not accuracy (*Chisq*_(1)_= 0.002, p=0.969), such that larger reinforcers led to greater speeding but no adjustment in accuracy when expressed as positive reinforcement relative to when they were expressed as negative reinforcement.

### Motivational context-specific effort adjustments are consistent with predictions of a normative model of control allocation

We previously showed that engaging in distinct control strategies in response to reward and punishment is normative under the assumption that participants are choosing control configurations that maximize reward rate while minimizing effort costs (where costs are operationalized as larger increases to one’s rate of evidence accumulation) (Leng et al., 2021). Specifically, we showed that such a reward-rate (RR) optimizing model predicts, and findings confirm, that larger rewards for correct responses should engender increases in *drift rate* (cf. attentional control) and decreases in *threshold* (cf. caution), whereas larger penalties for incorrect response should primarily yield increases in threshold. When fitting behavioral data (accuracies and reaction times) from the Collector Game to a Hierarchical Drift Diffusion model (HDDM) (Wiecki et al., 2013), we replicate both of these patterns (Fig 3 B & C, ps<0.001, Table S6).

We previously applied this RR model only to a context in which correct responses were incentivized by rewards (as in the Collector Game). However, it can be shown that the model’s predictions generalize to contexts in which correct responses are incentivized by aversive outcomes (Fig 3A ; see also Yee et al., 2022). That is, the RR model predicts that regardless of whether correct responses are incentivized by positive incentives (e.g., rewards) or negative incentives (e.g., loss avoidance), this should serve to reinforce behavior in the same manner (e.g., increased drift rate and decreased threshold) (Fig 3A). Importantly, this reinforcement-related enhancement of attentional control should be distinct from punishment-related enhancement of response caution, which should yield increased decision thresholds irrespective of the motivational context. Confirming these predictions, we found that negative incentives in the Protector Game produced distinct influences on DDM parameters depending on whether they were attached to correct responses (with greater loss avoidance resulting in higher drift rates p=0.01, and lower thresholds p<0.001) or incorrect responses (with higher penalties resulting in higher thresholds, p<0.001, and no significant change in drift rate, p=0.37) (Fig 3B and Table S7). In other words, the effects of reinforcement versus penalty on DDM parameters in the Protector Game exactly mirrored those in the Collector Game.

We previously showed that this same model can be applied, in reverse, to infer a given person’s weighting of rewards and punishments based only on their patterns of behavior in a given incentive condition (i.e., blind to the actual levels of reward and punishment in that condition). These reward and punishment estimates of weights were referred to as 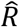 and 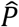, respectively. By extending the model to conditions of loss avoidance for a correct response (in our Protector Game), we should expect 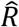 to generalize to estimates of any kind of reinforcement (positive or negative) and to therefore find similar variability in 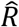 across levels of negative reinforcement as previously observed for positive reinforcement, and for these estimates to vary distinctly from 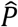 estimates.

Applying this inverse optimization approach to our current study, we replicate the model’s ability to infer higher values for higher incentive magnitude (*F*_(1,723)_ = 9.05, p=0.003) with higher 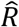 values for higher levels of reward (t(90)=10.83, p< 0.001), and higher 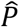 values for higher levels of punishment (t(90)=11.70, p< 0.001), in our Collector Game (Fig 4D). Critically, when applying this same approach to our novel Protector Game, we replicate this incentive magnitude effect (*F*_(1,723)_ = 6.92, p=0.009) whereby 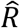 values are inferred to be higher with higher loss avoidance (t(90)=6.21, p<0.001), despite these being in the aversive rather than appetitive domain. 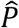 values were again higher for higher levels of punishment (t(90)=9.77, p<0.001). We also find that 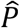 estimates were higher than 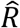 estimates in both the Collector game (*F*_(1,723)_= 1974.62, p<0.001) replicating previous findings, but we also found this same differentiation in the Protector Game,(*F*_(1,723)_= 1299.89, p<0.001) despite both being in the aversive domain. Concatenating estimates from both games we find that the main effect of incentive magnitude remained (F_(1,1447)_ = 15.83, p <0.001), 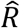 values were estimated to be higher for higher levels of reinforcement (t(181) =11.06, p <0.001) and 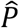 values were estimated to be higher for higher levels of punishment (t(181) =15.12, p <0.001) Table S6). Overall, these findings confirm that distinct strategies are deployed given the motivational context of an outcome, such that expected negative reinforcement and negative penalty differential influence the adjustment of the combination of drift rate parameters in order to maximize the potential reward rate during the turn.

**Figure 3.**
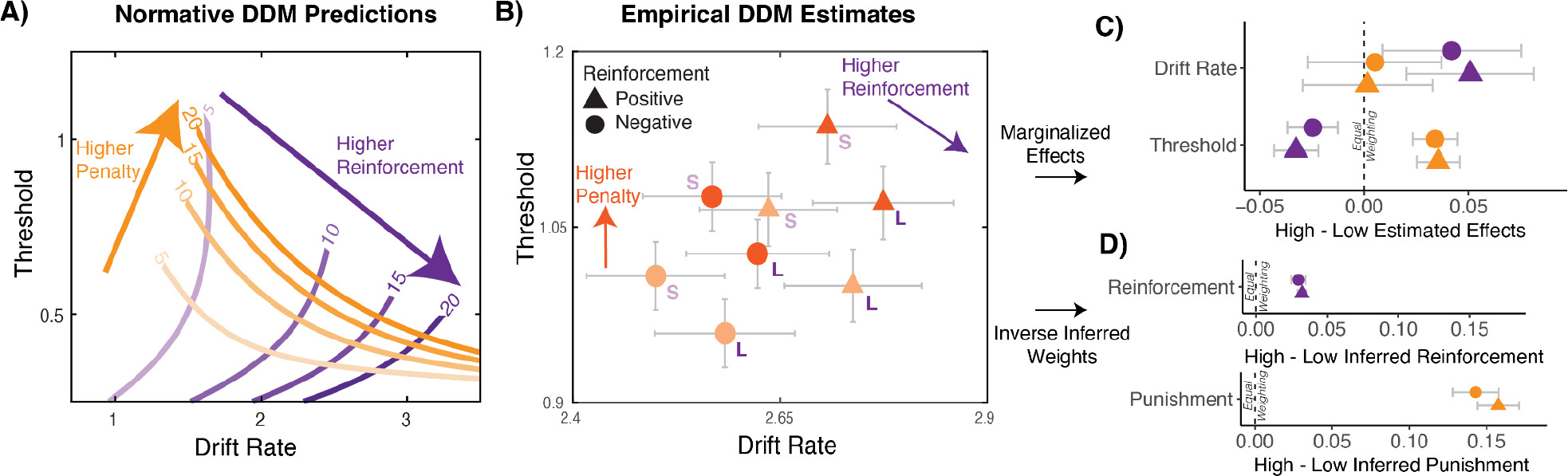
Normative and empirically observed estimates of incentive effects on drift diffusion model (DDM) parameters. (A-C). The normative model predicts that individuals select the combinations of drift rate and threshold that optimize (cost-discounted) reward rate, under different values of reinforcement and penalty. A key hypothesis of the model is that there is a clear dissociation between the extent to which higher levels of reinforcement are primarily associated with higher drift rate values versus how higher levels of penalty are primarily associated with higher threshold values. B) We fit our behavioral data to different parameterizations of the DDM, with drift rate and threshold varying with reinforcement, penalty, and/or congruence levels. Consistent with predictions of our reward rate optimization model, larger expected reinforcement (both reward and loss avoidance) was associated with increased drift rate and decreased threshold, whereas larger expected penalties led to increased threshold. Error bars reflect 95% Credible Intervals. C) Marginalized effects of drift rates and thresholds across all four experimental conditions (e.g., high minus low reinforcement marginalized across penalty magnitude. Error bars reflect s.d. D) Inverse inferred weights are calculated by taking participants’ estimated drift rate and thresholds for each condition then using the reward rate model to infer the reinforcement and penalty weights that best account for the individual’s behavior in that condition. Consistent with our previous finding from Leng et al., (2021), we demonstrate how inferred weights for high-low reinforcement is associated with task performance differences between high-low reinforcement, and higher sensitivity to penalty for the high vs. low penalty condition. Summary of individual-level contrasts between sensitivity to high vs. low reward and penalty. Error bars reflect s.e.m ***: p<0.001

**Figure 4.**
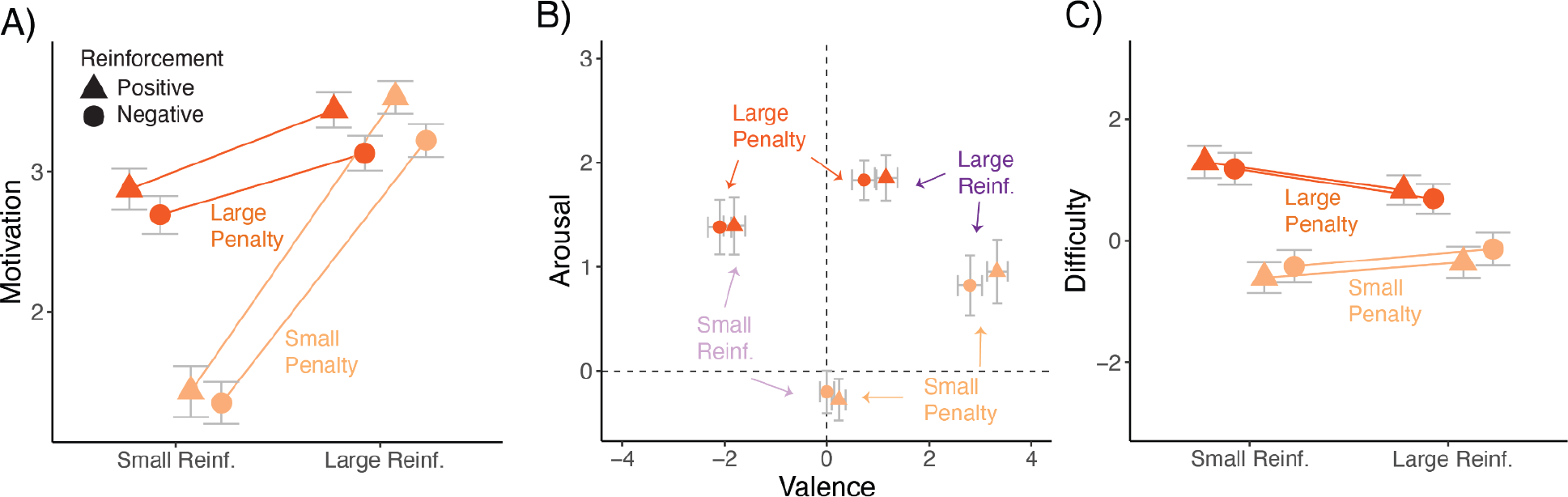
Subjective experience of composite motivation, valence, arousal, and difficulty by each incentive condition and game (Collector and Protector). A) Participants reported being more motivated by larger reinforcement and larger penalties. We observed a significant interaction indicating that participants were least motivated by the low reinforcement-low penalty condition compared to the other three conditions. B) Valence: Participants rated how positive or negative they found each of the conditions. We found that participants found larger reinforcements to be significantly more positive than smaller reinforcements and larger penalties to be significantly more negative than smaller penalties. Overall participants found the Collector Game to be more positive than the Protector Game. Arousal: Participants rated how excited to calm they found each of the conditions. We found that participants were more excited for both larger reinforcements and penalties compared to smaller reinforcements and penalties. C) Subjective experience of difficulty in each incentive condition in the Collector and Protector Game. Overall, we found that participants found conditions with larger penalties but not larger reinforcement more difficult. There were no interactions and this pattern was the same across the two games. Error bars reflect standard errors.

### Incentive types and ensuing effort adjustments give rise to distinct patterns of subjective experience

We show that behavior varies reliably depending on the type and magnitude of incentive, and account for this variability with a model that assumes that these incentives induce changes in motivational state and corresponding adjustments in control. However, neither in these experiments nor in our previous work (Leng et al., 2021) have we so far examined *experienced* motivational states themselves. It is therefore unclear to what extent participants experience this task as motivationally demanding and, to the extent they do, to what extent these experiences reflect variability in incentive type (reinforcement vs. punishment), incentive magnitude (small vs. large), and/or the magnitude and type of control adjustment (drift rate, threshold).To better understand how motivational state varied across our experimental conditions, at the end of the experiment we showed participants the cues for each condition across the games and asked them to rate their experience of effort, attention, and motivation during the associated intervals. We found that motivation, effort, and attention ratings were positively correlated with one another (Study 1 Collector: rs>.4, p<0.001; Study 1 Protector, rs>.4, p<0.001; Fig. S10) and therefore averaged them together to form a composite index of motivation (for analyses of each rating separately, see Table S8, S9, S10).

When examining the influence of explicit incentives on composite ratings of motivation, we found that in both the Collector and Protector games participants were most motivated by larger reinforcements (Collector: t(358)= 6.72, p<0.001; Protector: t(358)=5.90, p<0.001) and larger penalties (Collector: t(358)= 4.05, p<0.001; Protector: t(358)=3.73, p<0.001)(Fig 4A, Table S11, S12). We also observed an interaction such that participants were the least motivated when reinforcements and penalties were small (Collector: t(358)= -4.30, p<0.001; Protector: t(358)= - 3.90, p<0.001) (Fig 4A, Table S11,S12). The game type did not interact with the motivation reported for reinforcement (t(718)= 0.31, p=0.76) or penalty magnitude (t(718)= 0.07, p=0.94) (Table S13). Notably, this pattern did not directly mirror adjustments in either drift rate or threshold, suggesting that they did not reflect a direct readout of incentive magnitude (cf. Fig 1), behavior (cf. Fig 2AB and S9), or control adjustments (cf. Fig 4B) on their own. This suggests that these motivation ratings might reflect the alignment between a given incentive and the changes in control adjustment it induces, consistent with our model-based predictions and findings (cf. Fig. 3).

When examining subjective reports of valence, arousal, and difficulty, we found two patterns of findings that were distinct from what we observed for ratings of motivation. First, perhaps unsurprisingly, we find that across games participants rate higher levels of reinforcement (larger rewards in the Collector Game, larger loss avoided in the Protector Game) as more pleasant (Collector: t(358)= 17.53, p<0.001; Protector t(358)=15.95, p<0.001) and larger penalties as less pleasant (Collector: t(358)=-13.55, p<0.001; Protector: t(358)=-12.83, p<0.001) (Fig 5A, Table S8 & S9). Participants also found the Collector Game to be more pleasant than the Protector Game (t(718)= 3.12, p=0.002) (Table S10). No interactions emerged for valence ratings. Interestingly, while reinforcement and punishment induced opposite valences of emotional experience, we found that higher levels of *either* anticipated reinforcement (Collector: t(358)= 3.33, p<0.001; Protector t(358)=2.98, p=0.003) or punishment (Collector: t(358)= 4.71, p<0.001; Protector t(358)=5.07, p<0.001) were rated as more arousing than lower levels (Fig 5A, Table S8 & S9). The amount of arousal induced by these respective conditions did not differ between the two games (t(718)= 0.15, p=0.88) (Table S10).

A second pattern of results emerged unexpectedly. Despite task difficulty (i.e., average levels of response congruency) being held constant across our incentive conditions, we found that participants experienced intervals with higher penalties as being more difficult (Fig 4C, Table S8,S9,S10 Collector: t(358)= 6.05, p<0.001; Protector: t(358)= 4.58, p<0.001). This effect was selective to punishment levels – with no significant effects of reinforcement level (Collector: t(358)= -0.42 p= 0.675; Protector: t(358)=-0.36, p=0.716). We did not find that there was a difference in perceptions of difficulty based on whether the game was Collector or Protector(t(718)= -0.24, p=0.807, (Table S8)). We found a marginally significant interaction between reinforcement and punishment level (t(718)=-2.13, p=0.033), whereby penalty-related increases in difficulty ratings were numerically smaller with larger reinforcements.

## Discussion

Whether we are studying to ace an exam or preparing a paper for publication, succeeding requires summoning the motivation to exert the effort necessary. The motivation we bring to bear on the task is not only multiply determined (e.g., influenced by the consequences of achievement and failure), but it can also determine multiple changes in our actions and control strategies (e.g., whether we are more vigorous or cautious when exerting mental effort). We are not merely driven to put more effort into studying, but to study in particular ways (e.g., cramming or interleaving). Similarly, we may direct our paper-writing efforts toward efficiency or accuracy. The links between the multiple determinants and manifestations of effort remain poorly understood, including in particular the extent to which outcome valence forms a common link between these (e.g., with positive outcomes engendering different types of effort than aversive outcomes). Our findings show that the influence of a given outcome valence on the distribution of effort depends heavily on the motivational context for those outcomes (i.e., whether they serve as reinforcement or punishment).

Our findings build on past work demonstrating that people exert different types of effort for allocating cognitive control based on the expected positive outcomes for good performance and negative outcomes for poor performance (Leng et al., 2021). Using our novel Multi-Incentive Control Task which orthogonalizes incentive valence (positive vs. negative) and motivational context (reinforcement vs. punishment), we found that motivational context dominates valence in determining the influence of a given incentive (in addition to incentive magnitude). Specifically, patterns of behavior when losses were (inversely) contingent on correct responses (negative reinforcement) were entirely distinct from patterns observed when losses were instead contingent on incorrect responses (punishment). The former pattern closely mirrored patterns observed when correct responses were instead tied to reward gains (positive reinforcement). We showed that these findings were collectively well accounted for by a model that assumes participants configure cognitive control in a way that maximizes expected reward rate (cf. Bogacz et al., 2010; Leng et al., 2021; Shenhav et al., 2013). Together, these findings provide a potential novel lens to understand the past heterogeneity of findings for how aversive outcomes influence cognitive control adjustments (Braem et al., 2013; Cubillo et al., 2019; Levy & Schiller, 2021; Mobbs et al., 2020; Yee et al., 2015, 2021).

We found that these incentives also promoted distinct subjective experiences. Larger reinforcement led to greater positive arousal whereas larger punishment led to greater negative arousal. The effects of incentive type and magnitude on valence and arousal were the same regardless of motivational context, though participants did rate the positive reinforcement context (Collector game) as being more pleasant overall than the negative reinforcement context (Protector game). By contrast, the level of motivation participants felt (indexed by a composite of motivation, effort, and attention ratings) increased with higher levels of either reinforcement or punishment. Interestingly, these overall motivation ratings were not a direct read-out of a single type of control signal that was being adjusted (either drift rate or threshold; Fig. S9), consistent with the idea that motivation reflects a multivariate configuration across relevant control signals. Future work should seek to map these subjective ratings to the resulting control adjustments, as has been done previously in the case of univariate adjustments (Saunders et al., 2015; Yee et al., 2021; Corlazzoli et al., 2023).

In both games, an unexpected and intriguing finding emerged whereby participants perceived conditions with larger punishment (but not reinforcement) as more difficult, despite task difficulty (i.e., average response congruency) being held constant across incentive conditions. It is possible that this reflects an inherent link (or heuristic) people use when judging the difficulty of a task, incorporating into this judgment the risks associated with making an error. That is, participants may have evaluated task demands based both on when errors were more likely to occur (e.g., as a function of task congruency, which was equated across conditions) and based on the consequences of those errors. This is consistent with past work suggesting that error likelihood informs emotional responses to cognitive tasks (e.g., Inzlicht et al., 2015; Yang et al., 2023) as well as more recent findings that show that errors (and perhaps the potential for negative feedback that comes with them) induce feelings of fatigue (Matthews et al., 2023), suggesting a tighter link between these sources of negative affect than previously noted.

When comparing effort allocation between games, we found that participants performed similarly when working to earn monetary rewards as when they were working to avoid monetary losses. Initially, this seems at odds with predictions from Prospect Theory, which argues that individuals should exert more effort for loss avoidance relative to gains (Kahneman & Tversky, 1979).

However, there are at least two potential explanations for the apparent absence of such a loss aversion effect in our data. First, because the games were independent (and thus cumulative gains or losses avoided did not carry over), there was no explicit comparison in earnings between gain and loss avoidance conditions (e.g., participants would either work towards maximizing a positive amount of gems or work towards minimizing an initial loss of -300 gems per turn). Second, our mixed block design which varied outcome types between turns likely attenuated the participants’ ability to set a concrete reference point. Future research could test whether cognitive control strategies are impacted by loss aversion by allowing for cumulative earnings across both games. Here, we would predict that while the motivational context would determine the control strategy used (e.g., reinforcement promoting increased drift rate, punishment promoting increased decision threshold), we might also observe that motivation to avoid losses would promote increased overall control allocation relative to gains.

Overall, these data contribute to a growing literature demonstrating that the effects of aversive motivation on cognitive control is context-dependent (Lindström et al., 2013; Millner et al., 2018; Yang et al., 2023). Our work highlights how computational approaches provide a more precise understanding of how people select diverse strategies for allocating cognitive control based on the type and context of a given motivational incentive. A promising future direction would be to test to what extent these control strategies apply to diverse types of aversive outcomes (e.g., shocks), which would provide a richer understanding of the generalizability of our model’s normative predictions of mental effort allocation across diverse incentives (Crawford et al., 2020; Kray et al., 2018). These findings further provide novel insight into how the congruency between expected outcomes and behavioral goals determine our actions (and computational strategies) in value-based decision-making tasks (De Martino & Cortese, 2023; Frömer et al., 2019; Guitart-Masip et al., 2014; Molinaro & Collins, 2023). Consistent with recent work on this topic, our data show that whether our actions are aligned with our goals matters as much (if not more) than the valence of the outcomes.

An important avenue for this work is its potential applicability to computational psychiatry, which aims to leverage neurocomputational mechanisms to elucidate the basis of disorder or symptom-specific impairments in clinical disorders (Huys et al., 2016; Montague et al., 2012; Yip et al., 2023). Delineating how these motivational factors drive different types of cognitive control provides a crucial intermediary step for informing how these processes are disrupted in mood and psychotic disorders (e.g., major depressive disorder and schizophrenia; (Barch et al., 2018; Culbreth et al., 2016; Grahek et al., 2019; Joormann & Vanderlind, 2014; Paulus, 2015), as well as how they might vary with experience (Hanson et al., 2017; Machlin et al., 2019; Sheridan & McLaughlin, 2014) and development (C. Geier & Luna, 2009; Insel et al., 2017; Paulsen et al., 2015; Somerville & Casey, 2010) within the broader population.

## Acknowledgments

This research was supported by an NSF GRFP Fellowship (MPF), an NIH T32-MH126388 (DY), NIH-R25-NS124530 (DY), an NIH T32-MH115895 (XL), an NSF CAREER Award 2046111 (AS), and an NIH Award 1S10OD025181 (AS). We would like to thank Carolyn K. Dean Wolf for her help designing the task along with all members of the Shenhav Lab for providing feedback at all stages of the project from design to writing.

## Author Contributions

The authors confirm contribution to the paper as follows: study conception and design: DY, XL, MPF, MT, AS; data collection: MPF; analysis and interpretation of results: AS, MPF, XL, DY; draft manuscript preparation: MPF, DY, AS. All authors reviewed the results and approved the final version of the manuscript.

## Data Availability

Data for the study will be made available upon publication.

## Code Availability

Analysis code will be made available upon publication.

### Preregistration

The study reported in this article was not preregistered.

