## Supplemental for "Motivational context determines the impact of aversive outcomes on mental effort allocation" (from another version of this preprint: version 1, 2023-11-01)

### Supplemental Material

#### Behavior

##### **Table S1.** Mixed Model results for Collector Game for Correct Response Per Second, Reaction Time, Accuracy

| **Collector** | | | | | | | | | | | | |
|---|---|---|---|---|---|---|---|---|---|---|---|---|
| | **Correct Response per Second** | | | | **Trial Accuracy** | | | | **Trial Logged Accurate Reaction Time** | | | |
| *Predictors* | *Estimates* | *CI* | *Statistic* | *p* | *Odds Ratios* | *CI* | *Statistic* | *p* | *Estimates* | *CI* | *Statistic* | *p* |
| Intercept | 1.07 | 0.99 – 1.14 | 27.83 | **<0.001** | 23.91 | 17.90 – 31.93 | 21.50 | **<0.001** | 2.90 | 2.88 – 2.92 | 254.13 | **<0.001** |
| Reinforcement Magnitude | 0.03 | 0.02 – 0.04 | 7.92 | **<0.001** | 0.93 | 0.88 – 1.00 | -2.06 | **0.039** | -0.01 | -0.01 – -0.01 | -9.19 | **<0.001** |
| Penalty Magnitude | -0.01 | -0.02 – -0.00 | -2.69 | **0.007** | 1.24 | 1.16 – 1.33 | 6.40 | **<0.001** | 0.01 | 0.01 – 0.01 | 7.01 | **<0.001** |
| Interval Congruency | -0.00 | -0.01 – 0.00 | -1.40 | 0.162 | | | | | | | | |
| Interval Length | 0.02 | 0.01 – 0.02 | 4.50 | **<0.001** | 1.00 | 0.95 – 1.05 | -0.02 | 0.983 | 0.01 | 0.01 – 0.01 | 9.07 | **<0.001** |
| Interval Number in Game | -0.01 | -0.02 – -0.00 | -2.81 | **0.005** | 0.95 | 0.90 – 0.99 | -2.21 | **0.027** | 0.00 | -0.00 – 0.00 | 1.41 | 0.158 |
| Order | 0.05 | -0.04 – 0.14 | 1.10 | 0.272 | 0.91 | 0.65 – 1.28 | -0.53 | 0.596 | -0.02 | -0.05 – 0.01 | -1.31 | 0.191 |
| Sex | 0.01 | -0.04 – 0.05 | 0.24 | 0.807 | 0.98 | 0.83 – 1.15 | -0.26 | 0.796 | -0.00 | -0.01 – 0.01 | -0.23 | 0.815 |
| Age | -0.02 | -0.06 – 0.02 | -0.98 | 0.326 | 1.23 | 1.04 – 1.45 | 2.39 | **0.017** | 0.01 | -0.00 – 0.02 | 1.55 | 0.122 |
| Reinf. Magnitude x Penalty Magnitude | 0.01 | -0.00 – 0.01 | 1.54 | 0.124 | 0.97 | 0.92 – 1.02 | -1.30 | 0.194 | -0.00 | -0.00 – 0.00 | -1.52 | 0.129 |
| Trial Congruency | | | | | 1.13 | 1.04 – 1.24 | 2.69 | **0.007** | -0.05 | -0.05 – -0.05 | -32.69 | **<0.001** |
| Trial Number in Interval | | | | | 0.75 | 0.72 – 0.79 | -12.12 | **<0.001** | -0.04 | -0.04 – -0.04 | -47.66 | **<0.001** |
| **Random Effects** | | | | | | | | | | | | |
| $\sigma^2$ | 0.06 | | | | 3.29 | | | | 0.02 | | | |
| $\tau_{00}$ | 0.04 SubID | | | | 0.54 SubID | | | | 0.00 SubID | | | |
| $\tau_{11}$ | 0.00 SubID.ValueLevel1 | | | | 0.02 SubID.ValueLevel1 | | | | 0.00 SubID.ValueLevel1 | | | |
| | 0.00 SubID.PenaltyLevel1 | | | | 0.02 SubID.PenaltyLevel1 | | | | 0.00 SubID.PenaltyLevel1 | | | |
| $\rho_{01}$ | -0.03 | | | | 0.39 | | | | -0.04 | | | |
| | 0.13 | | | | -0.32 | | | | -0.06 | | | |
| ICC | 0.40 | | | | 0.15 | | | | 0.17 | | | |

| | | | |
|---|---|---|---|
| N | 91 SubID | 91 SubID | 91 SubID |
| Observations | 5412 | 35987 | 33974 |
| Marginal $R^2$ / Conditional $R^2$ | 0.024 / 0.416 | 0.048 / 0.192 | 0.091 / 0.249 |

**Table S2.** Mixed Model results for Protector Game for Correct Response Per Second, Reaction Time, Accuracy

| **Protector** | | | | | | | | | | | | |
|---|---|---|---|---|---|---|---|---|---|---|---|---|
| | **Correct Response per Second** | | | | **Trial Accuracy** | | | | **Trial Logged Accurate Reaction Time** | | | |
| *Predictors* | *Estimates* | *CI* | *Statistic* | *p* | *Odds Ratios* | *CI* | *Statistic* | *p* | *Estimates* | *CI* | *Statistic* | *p* |
| Intercept | 1.10 | 1.02 – 1.18 | 27.68 | **<0.001** | 21.26 | 15.06 – 30.01 | 17.39 | **<0.001** | 2.88 | 2.86 – 2.91 | 247.63 | **<0.001** |
| Reinforcement Magnitude | 0.03 | 0.02 – 0.04 | 5.28 | **<0.001** | 0.92 | 0.84 – 1.01 | -1.83 | 0.068 | -0.01 | -0.01 – -0.01 | -5.84 | **<0.001** |
| Penalty Magnitude | -0.01 | -0.02 – -0.00 | -2.32 | **0.020** | 1.17 | 1.07 – 1.27 | 3.61 | **<0.001** | 0.01 | 0.01 – 0.01 | 5.99 | **<0.001** |
| Interval Congruency | -0.00 | -0.01 – 0.00 | -1.36 | 0.174 | | | | | | | | |
| Interval Length | 0.01 | 0.00 – 0.02 | 2.50 | **0.012** | 1.00 | 0.95 – 1.04 | -0.08 | 0.935 | 0.01 | 0.01 – 0.01 | 10.32 | **<0.001** |
| Interval Number in Game | 0.01 | 0.00 – 0.02 | 2.05 | **0.040** | 1.05 | 1.00 – 1.11 | 1.98 | **0.048** | -0.00 | -0.00 – -0.00 | -2.34 | **0.019** |
| Order | -0.01 | -0.10 – 0.08 | -0.18 | 0.857 | 0.99 | 0.66 – 1.49 | -0.05 | 0.961 | 0.00 | -0.02 – 0.03 | 0.34 | 0.736 |
| Sex | 0.03 | -0.02 – 0.07 | 1.15 | 0.248 | 1.00 | 0.83 – 1.22 | 0.04 | 0.965 | -0.00 | -0.02 – 0.01 | -0.63 | 0.526 |
| Age | -0.04 | -0.08 – 0.01 | -1.63 | 0.102 | 0.99 | 0.81 – 1.19 | -0.15 | 0.877 | 0.01 | -0.00 – 0.02 | 1.61 | 0.106 |
| Reinf. Magnitude x Penalty Magnitude | -0.00 | -0.01 – 0.00 | -0.78 | 0.435 | 0.94 | 0.89 – 0.99 | -2.57 | **0.010** | 0.00 | -0.00 – 0.00 | 0.24 | 0.807 |
| Trial Congruency | | | | | 1.32 | 1.20 – 1.44 | 5.93 | **<0.001** | -0.05 | -0.05 – -0.05 | -33.20 | **<0.001** |
| Trial Number in Interval | | | | | 0.80 | 0.77 – 0.84 | -9.44 | **<0.001** | -0.04 | -0.04 – -0.03 | -46.03 | **<0.001** |
| **Random Effects** | | | | | | | | | | | | |
| $\sigma^2$ | 0.07 | | | | 3.29 | | | | 0.02 | | | |
| $\tau_{00}$ | 0.05 SubID | | | | 0.86 SubID | | | | 0.00 SubID | | | |
| $\tau_{11}$ | 0.00 SubID.ValueLevel1 | | | | 0.10 SubID.ValueLevel1 | | | | 0.00 SubID.ValueLevel1 | | | |
| | 0.00 SubID.PenaltyLevel1 | | | | 0.09 SubID.PenaltyLevel1 | | | | 0.00 SubID.PenaltyLevel1 | | | |
| $\rho_{01}$ | -0.28 | | | | -0.22 | | | | -0.00 | | | |
| | 0.33 | | | | -0.32 | | | | 0.15 | | | |
| ICC | 0.44 | | | | 0.24 | | | | 0.19 | | | |
| N | 91 SubID | | | | 91 SubID | | | | 91 SubID | | | |
| Observations | 5433 | | | | 36194 | | | | 33872 | | | |
| Marginal $R^2$ / Conditional $R^2$ | 0.024 / 0.452 | | | | 0.024 / 0.261 | | | | 0.084 / 0.256 | | | |

#### **Table S3.** Mixed Model results for Protector Game for Correct Response Per Second, Reaction Time, Accuracy

| | Correct Response per Second | | | | Trial Accuracy | | | | Trial Logged Accurate Reaction Time | | | |
|---|---|---|---|---|---|---|---|---|---|---|---|---|
| *Predictors* | *Estimates* | *CI* | *Statistic* | *p* | *Odds Ratios* | *CI* | *Statistic* | *p* | *Estimates* | *CI* | *Statistic* | *p* |
| Intercept | 1.08 | 1.01 – 1.16 | 28.53 | **<0.001** | 22.77 | 17.10 – 30.30 | 21.42 | **<0.001** | 2.89 | 2.87 – 2.91 | 256.40 | **<0.001** |
| Reinforcement Magnitude | 0.03 | 0.02 – 0.04 | 8.93 | **<0.001** | 0.94 | 0.88 – 0.99 | -2.14 | **0.033** | -0.01 | -0.01 – -0.01 | -9.19 | **<0.001** |
| Penalty Magnitude | -0.01 | -0.02 – -0.00 | -3.19 | **0.001** | 1.20 | 1.13 – 1.27 | 6.28 | **<0.001** | 0.01 | 0.01 – 0.01 | 7.99 | **<0.001** |
| Reinforcement Valence | -0.02 | -0.04 – 0.00 | -1.56 | 0.120 | 1.04 | 0.89 – 1.21 | 0.49 | 0.623 | 0.01 | 0.00 – 0.01 | 2.88 | **0.004** |
| Interval Congruency | -0.00 | -0.01 – 0.00 | -1.69 | 0.090 | | | | | | | | |
| Interval Length | 0.01 | 0.01 – 0.02 | 4.92 | **<0.001** | 1.00 | 0.96 – 1.03 | -0.17 | 0.866 | 0.01 | 0.01 – 0.01 | 13.74 | **<0.001** |
| Interval Number in Session | -0.00 | -0.01 – 0.01 | -0.34 | 0.736 | 1.00 | 0.93 – 1.07 | 0.02 | 0.980 | -0.00 | -0.00 – 0.00 | -0.68 | 0.496 |
| Order | 0.02 | -0.07 – 0.11 | 0.52 | 0.602 | 0.92 | 0.66 – 1.30 | -0.45 | 0.651 | -0.01 | -0.03 – 0.02 | -0.52 | 0.601 |
| Sex | 0.02 | -0.02 – 0.06 | 0.95 | 0.343 | 1.00 | 0.86 – 1.16 | -0.00 | 0.996 | -0.00 | -0.02 – 0.01 | -0.47 | 0.637 |
| Age | -0.03 | -0.07 – 0.01 | -1.33 | 0.184 | 1.15 | 0.98 – 1.35 | 1.76 | 0.078 | 0.01 | -0.00 – 0.02 | 1.62 | 0.105 |
| Reinf. Magnitude x Penalty Magnitude | 0.00 | -0.00 – 0.01 | 0.51 | 0.611 | 0.96 | 0.93 – 0.99 | -2.43 | **0.015** | -0.00 | -0.00 – 0.00 | -0.89 | 0.374 |
| Reinf. Magnitude x Reinf. Valence | 0.00 | -0.00 – 0.01 | 1.06 | 0.291 | 1.00 | 0.97 – 1.03 | -0.04 | 0.969 | -0.00 | -0.00 – -0.00 | -2.59 | **0.010** |
| Penalty x Reinf. Valence | 0.00 | -0.00 – 0.01 | 0.15 | 0.877 | 1.01 | 0.98 – 1.05 | 0.55 | 0.580 | 0.00 | -0.00 – 0.00 | 0.34 | 0.736 |
| Reinf. Valence x Order | 0.03 | 0.00 – 0.06 | 1.96 | **0.050** | 0.97 | 0.79 – 1.18 | -0.31 | 0.755 | -0.01 | -0.02 – -0.00 | -2.85 | **0.004** |
| Reinf. Magnitude x Penalty Magnitude x Reinf. Valence | 0.00 | -0.00 – 0.01 | 1.65 | 0.100 | 1.01 | 0.97 – 1.04 | 0.39 | 0.695 | -0.00 | -0.00 – 0.00 | -1.17 | 0.241 |
| Trial Congruencey | | | | | 1.22 | 1.15 – 1.30 | 6.11 | **<0.001** | -0.05 | -0.05 – -0.05 | -46.55 | **<0.001** |
| Trial Number in Interval | | | | | 0.78 | 0.75 – 0.80 | -15.17 | **<0.001** | -0.04 | -0.04 – -0.04 | -66.44 | **<0.001** |
| **Random Effects** | | | | | | | | | | | | |
| $\sigma^2$ | 0.07 | | | | 3.29 | | | | 0.02 | | | |
| $\tau_{00}$ | 0.04 SubID | | | | 0.60 SubID | | | | 0.00 SubID | | | |
| $\tau_{11}$ | 0.00 SubID.ValueLevel1 | | | | 0.05 SubID.ValueLevel1 | | | | 0.00 SubID.ValueLevel1 | | | |
| | 0.00 SubID.PenaltyLevel1 | | | | 0.04 SubID.PenaltyLevel1 | | | | 0.00 SubID.PenaltyLevel1 | | | |
| | 0.00 SubID.ValenceLevel1 | | | | 0.12 SubID.ValenceLevel1 | | | | 0.00 SubID.ValenceLevel1 | | | |
| $\rho_{01}$ | -0.18 | | | | 0.06 | | | | -0.01 | | | |

| | | | |
|---|---|---|---|
| | 0.29 | -0.45 | 0.03 |
| | -0.16 | -0.31 | -0.04 |
| ICC | 0.41 | 0.20 | 0.18 |
| N | 91 SubID | 91 SubID | 91 SubID |
| Observations | 10845 | 72181 | 67846 |
| Marginal $R^2$ / Conditional $R^2$ | 0.023 / 0.428 | 0.033 / 0.225 | 0.088 / 0.250 |

#### Figure S1. Correct Response Per Second for Collector and Protector Games.

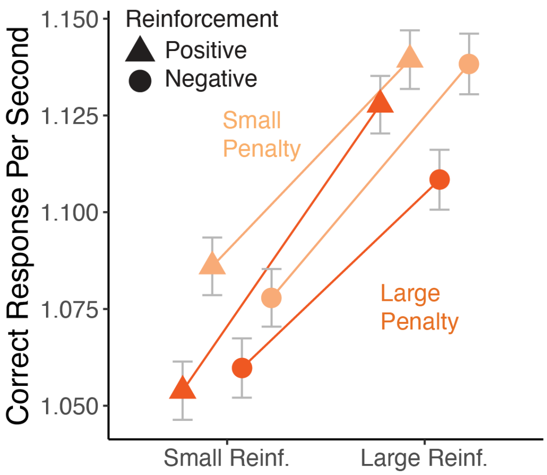


**Figure S1. Correct Response Per Second for Collector and Protector Games.** Effects of aversive and positive reinforcement and punishment on performance during turns. A) With increasing expected reinforcement in the positive context participants completed more trials correctly per second for higher reinforcement whereas they completed fewer trials correctly per second when they expected higher penalties in the turn. B) Expectations of higher penalties in the aversive context similarly promoted completing fewer trials correctly per second. In contrast, when expecting higher negative reinforcement participants completed more trials correctly per second. Error bars reflect standard errors.

### HDDM

#### **Table S4.** List of Models

| Which data models were run on | Formula | Intertrial-variability |
|---|---|---|
| Collector,<br>Protector,<br>Both Games Together | a~Reinf.Magnitude+Pen.Magnitude',<br>'v~Reinf.Magnitude+Pen.Magnitude+congruency',<br>'z~congruency' | p_outlier |
| Collector,<br>Protector,<br>Both Games Together | a~Reinf.Magnitude+Pen.Magnitude+ scaledRunningTime',<br>'v~Reinf.Magnitude+Pen.Magnitude+congruency',<br>'z~congruency' | p_outlier', 'sv', 'st' |
| Collector,<br>Protector,<br>Both Games Together | a~Reinf.Magnitude*Pen.Magnitude+ scaledRunningTime',<br>'v~Reinf.Magnitude*Pen.Magnitude+congruency',<br>'z~congruency' | p_outlier', 'sv', 'st' |
| Collector,<br>Protector,<br>Both Games Together | a~Reinf.Magnitude+Pen.Magnitude*scaledRunningTime',<br>'v~Reinf.Magnitude+Pen.Magnitude+congruency',<br>'z~congruency' | p_outlier', 'sv', 'st' |
| Collector,<br>Protector,<br>Both Games Together | a~Reinf.Magnitude*Pen.Magnitude + Pen.Magnitude*scaledRunningTime',<br>'v~Reinf.Magnitude*Pen.Magnitude+congruency',<br>'z~congruency' | p_outlier', 'sv', 'st' |
| Collector,<br>Protector,<br>Both Games Together | a~Reinf.Magnitude*Pen.Magnitude',<br>v~Reinf.Magnitude*Pen.Magnitude+congruency',<br>'z~congruency' | p_outlier |
| Collector,<br>Protector,<br>Both Games Together | a~Reinf.Magnitude*Pen.Magnitude',<br>'v~Reinf.Magnitude*Pen.Magnitude+congruency' | p_outlier', 'sv', 'st' |
| Collector,<br>Protector,<br>Both Games Together | a~Reinf.Magnitude*Pen.Magnitude',<br>'v~Reinf.Magnitude*Pen.Magnitude+congruency',<br>'z~congruency' | p_outlier', 'sv', 'st' |
| Collector,<br>Protector,<br>Both Games Together | a~Reinf.Magnitude+Pen.Magnitude',<br>'v~Reinf.Magnitude+Pen.Magnitude+congruency',<br>'z~congruency' | p_outlier', 'sv', 'st' |
| Collector,<br>Protector,<br>Both Games Together | a~Reinf.Magnitude+Pen.Magnitude',<br>'v~Reinf.Magnitude+Pen.Magnitude+congruency' | p_outlier |
| Collector,<br>Protector,<br>Both Games Together | a~Reinf.Magnitude+Pen.Magnitude',<br>'v~Reinf.Magnitude+Pen.Magnitude+congruency' | p_outlier', 'sv', 'st' |
| Collector,<br>Protector,<br>Both Games Together | a~Reinf.Magnitude*Pen.Magnitude',<br>'v~Reinf.Magnitude*Pen.Magnitude+congruency' | p_outlier |
| Study1_Both | a~Reinf.Magnitude*Reinf.Valence+Pen.Magnitude*Reinf.Valence+scaledRunningTime',<br>'v~Reinf.Magnitude*Reinf.Valence+Pen.Magnitude*Reinf.Valence+congruency',<br>'z~congruency' | p_outlier', 'sv', 'st' |
| Study1_Both | a~Reinf.Magnitude+Pen.Magnitude+Reinf.Valence+scaledRunningTime',<br>'v~Reinf.Magnitude+Pen.Magnitude+Reinf.Valence+congruency',<br>'z~congruency' | p_outlier', 'sv', 'st' |
| Study1_Both | a~Reinf.Magnitude+Pen.Magnitude+Reinf.Valence',<br>'v~Reinf.Magnitude+Pen.Magnitude+Reinf.Valence+congruency',<br>'z~congruency' | p_outlier', 'sv', 'st' |
| Study1_Both | a~Reinf.Magnitude*Reinf.Valence+Pen.Magnitude*Reinf.Valence',<br>'v~Reinf.Magnitude*Reinf.Valence+Pen.Magnitude*Reinf.Valence+congruency',<br>'z~congruency' | p_outlier', 'sv', 'st' |
| Study1_Both | a~Reinf.Magnitude*Pen.Magnitude+<br>Reinf.Magnitude*Reinf.Valence+Pen.Magnitude*Reinf.Valence+scaledRunningTime',<br>'v~Reinf.Magnitude*Pen.Magnitude+ Reinf.Magnitude*Reinf.Valence+Pen.Magnitude*Reinf.Valence+congruency',<br>'z~congruency' | p_outlier', 'sv', 'st' |
| Study1_Both | a~Reinf.Magnitude*Pen.Magnitude+Reinf.Magnitude*Reinf.Valence+Pen.Magnitude*Reinf.Valence+Pen.Magnitude*sc<br>aledRunningTime',<br>'v~Reinf.Magnitude*Pen.Magnitude+Reinf.Magnitude*Reinf.Valence+Pen.Magnitude*Reinf.Valence+congruency',<br>'z~congruency' | p_outlier', 'sv', 'st' |
| Study1_Both | a~Reinf.Magnitude+Pen.Magnitude+Order+scaledRunningTime',<br>'v~Reinf.Magnitude+Pen.Magnitude+Order+congruency',<br>'z~congruency' | p_outlier', 'sv', 'st' |
| Study1_Both | a~Reinf.Magnitude*Pen.Magnitude + Reinf.Magnitude*Order+Pen.Magnitude*Order+scaledRunningTime',<br>'v~Reinf.Magnitude*Pen.Magnitude+ Reinf.Magnitude*Order+Pen.Magnitude*Order+congruency',<br>'z~congruency' | p_outlier', 'sv', 'st' |

**Table S5.** Estimated Effect of Drift Rate and Threshold in Both Games Model

| Effect | Parameter | Mean (SD) | p value | sig | CI |
|---|---|---|---|---|---|
| Reinforcement *Large-Small* | Drift Rate | 0.05 (0.01) | 0.00 | *** | [0.02 0.07] |
| | Threshold | -0.03 (0.00) | 0.00 | *** | [-0.04 -0.02] |
| Penalty *Large-Small* | Drift Rate | 0.00 (0.01) | 0.44 | n.s. | [-0.02 0.03] |
| | Threshold | 0.03 (0.00) | 0.00 | *** | [0.03 0.04] |
| Valence *Positve-Negative* | Drift Rate | 0.01 (0.02) | 0.31 | n.s. | [-0.03 0.05] |
| | Threshold | 0.00 (0.01) | 0.32 | n.s. | [-0.01 0.02] |

Table S5 shows the details of the ANOVA of the effects of Reinforcement Magnitude, Penalty Magnitude, and Reinforcement Valence on Drift Rate and Threshold in the DDM Model that includes data from both the Collector and Protector Games.

**Table S6.** Estimated Effect of Drift Rate and Threshold in Collector only Model

| *Effect* | *Parameter* | *Mean (SD)* | *p value* | *sig* | *CI* |
|---|---|---|---|---|---|
| Reinforcement *Large-Small* | Drift Rate | 0.05 (0.02) | 0.00 | *** | [0.02 0.08] |
| | Threshold | -0.03 (0.01) | 0.00 | *** | [-0.04 -0.02] |
| Penalty *Large-Small* | Drift Rate | 0.00 (0.02) | 0.46 | n.s. | [-0.03 0.03] |
| | Threshold | 0.04 (0.01) | 0.00 | *** | [0.03 0.05] |

Table S6 shows the details of the anova of the effects of Reinforcement magnitude and Penalty Magnitude, on Drift Rate and Threshold in the Collector Game only DDM Model.

**Table. S7** Estimated Effect of Drift Rate and Threshold in Protector only Model

| *Effect* | *parameter* | *Mean (SD)* | *p value* | *sig* | *CI* |
|---|---|---|---|---|---|
| Reinforcement *Large-Small* | Drift Rate | 0.04 (0.02) | 0.01 | ** | [0.01 0.08] |
| | Threshold | -0.02 (0.01) | 0.00 | *** | [-0.04 -0.01] |
| Penalty *Large-Small* | Drift Rate | 0.01 (0.02) | 0.37 | n.s. | [-0.03 0.03] |
| | Threshold | 0.03 (0.01) | 0.00 | *** | [0.02 0.04] |

Table S7 shows the details of the anova of the effects of Reinforcement magnitude and Penalty Magnitude, on Drift Rate and Threshold from the HDDM model fit on reaction times and accuracies from the Protector Game only.

**Figure S2.** BPICs for Models

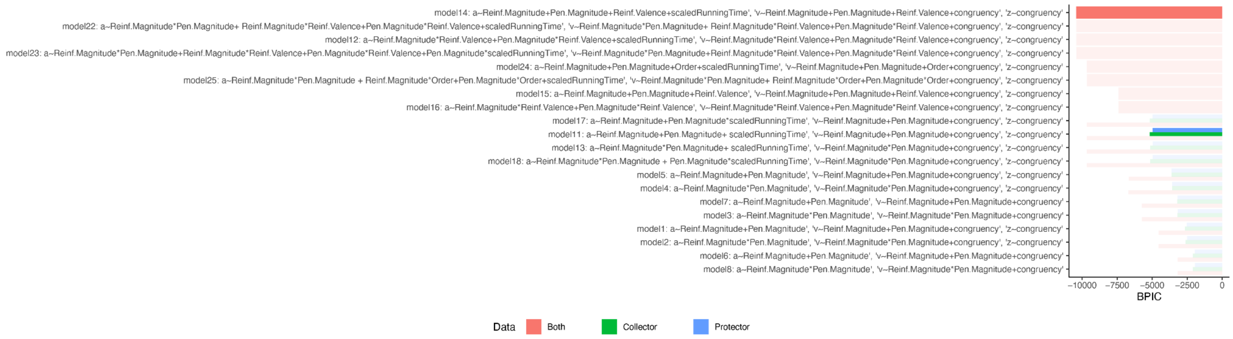


Figure S2. BPICs for HDDMs. Models were fit on either Collector Data, Protector Data or Both Collector and Protector Data. Fitting the models on data from both games in the same model allowed us to examine the influence of reinforcement valence on Drift and Threshold as well as game order. Winning Models as determined by the BPIC, Traces, and PPCs are highlighted.

**Figure S3.** Traces for Full Model

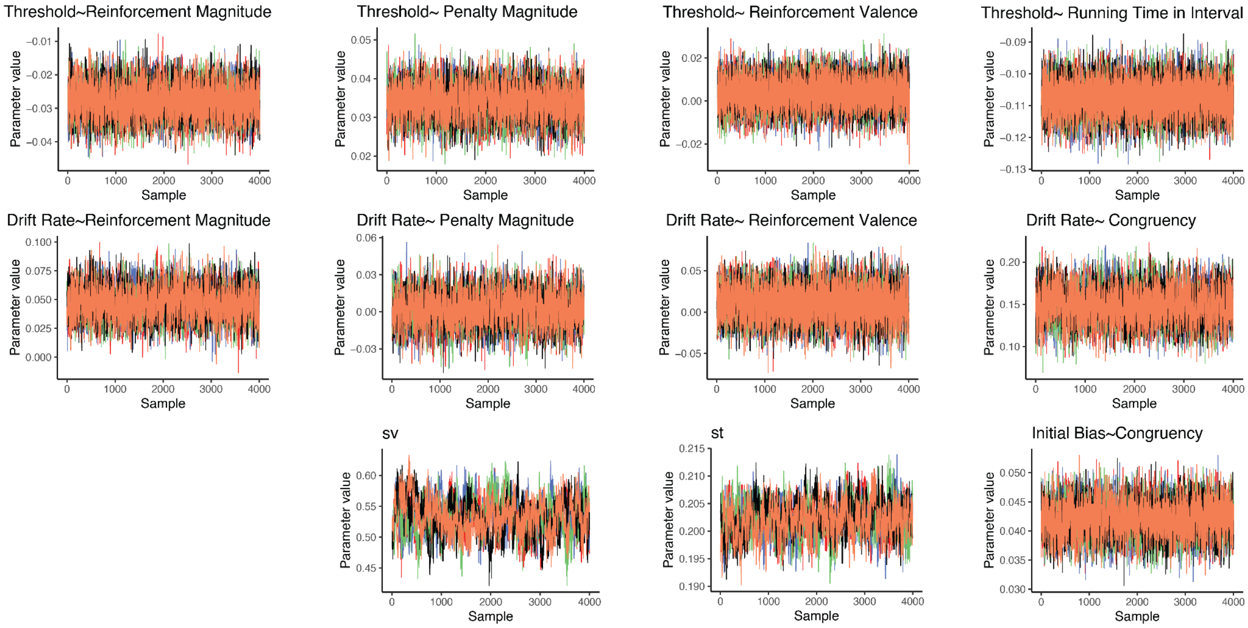


Figure S3. Trace Plots for Model fit on data from both Collector and Protector Games. Trace plots for the full model for each of the parameters. The trace plots visualize the parameter value selection and look like lovely fuzzy caterpillars indicating good convergence.

**Figure S4.** Posterior Predictive Check for Model fit on Data from Both Collector and Protector

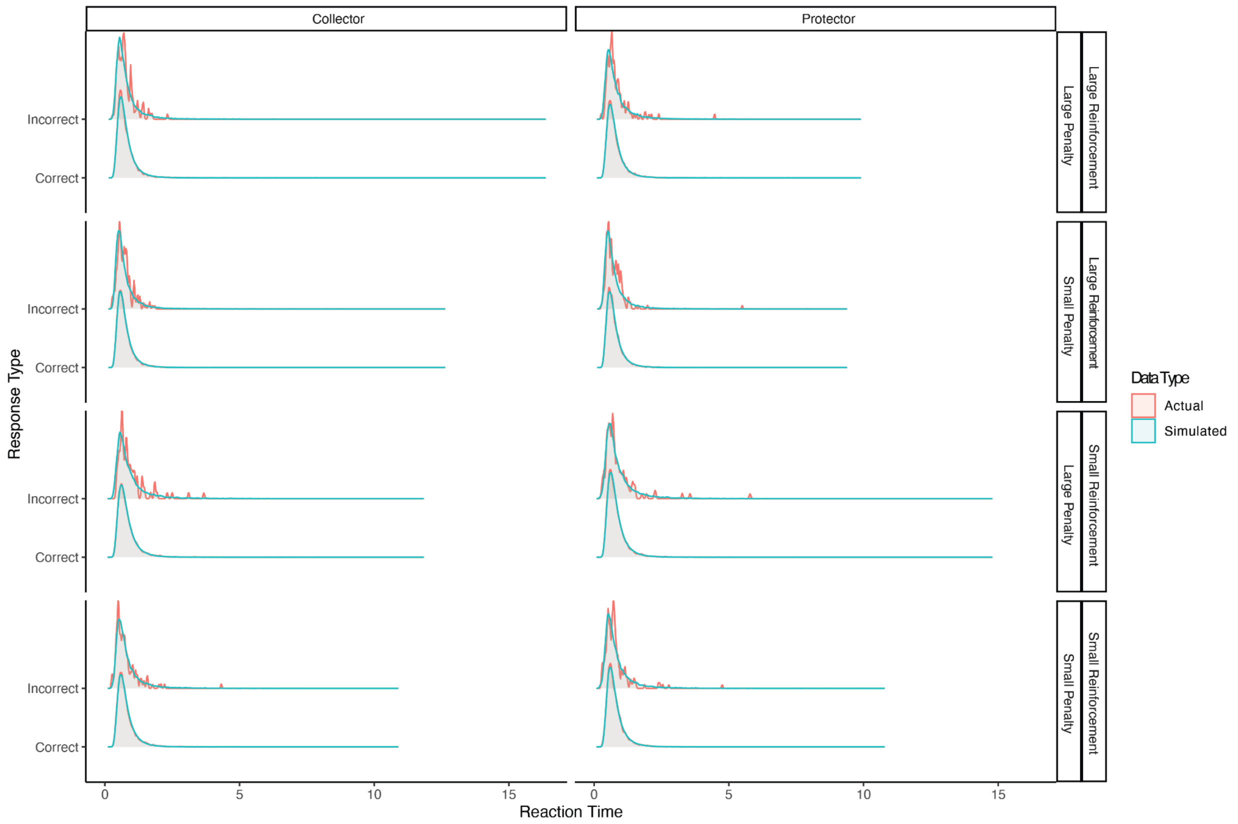


We performed a posterior predictive check for the fitted Drift Diffusion Model (DDM). We generated 500 independent samples from the posterior distribution of fitted parameters and then simulated the reaction time distribution for each posterior sample. The predicted reaction time distribution matches with the actual reaction time distribution for each condition.

**Figure S5.** Traces for Collector only Model

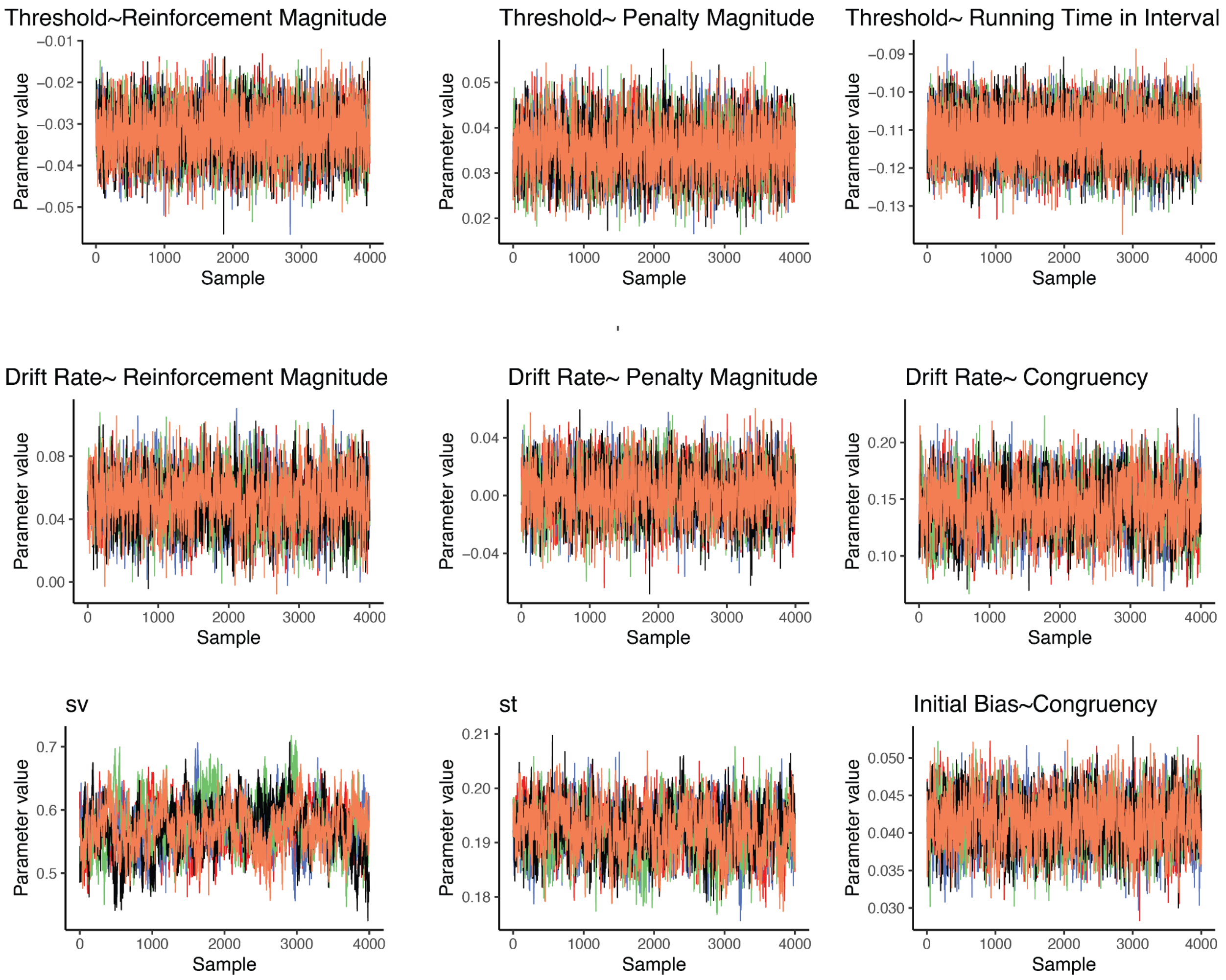


Figure 5 shows the trace plots for the Collector Game model only for each of the parameters. The Trace plots visualize the parameter value selection and look like lovely fuzzy caterpillars indicating good convergence.

**Figure S6.** Posterior Predictive Checks for Collector only Model

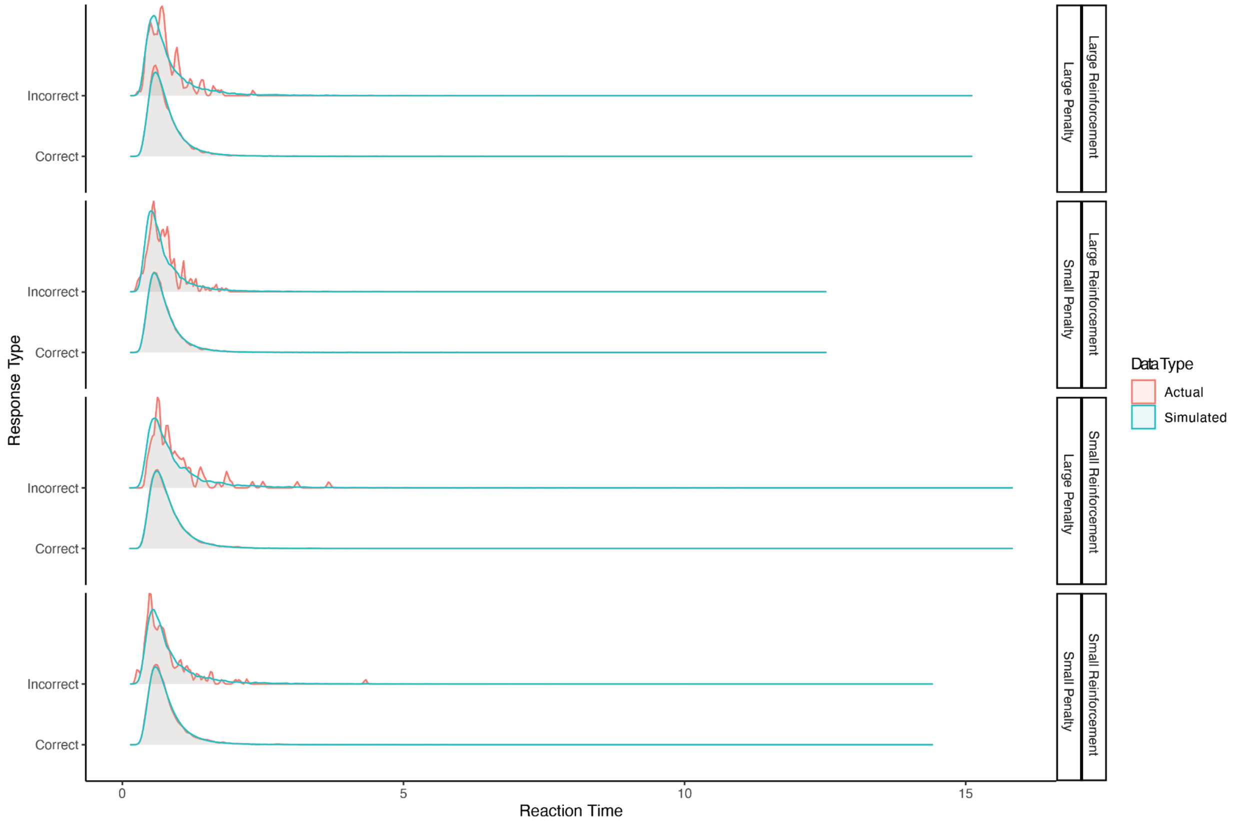


We performed a posterior predictive check for the fitted Drift Diffusion Model (DDM). We generated 500 independent samples from the posterior distribution of fitted parameters and then simulated the reaction time distribution for each posterior sample. The predicted reaction time distribution matches with the actual reaction time distribution for each condition.

**Figure S7.** Traces for Protector Model

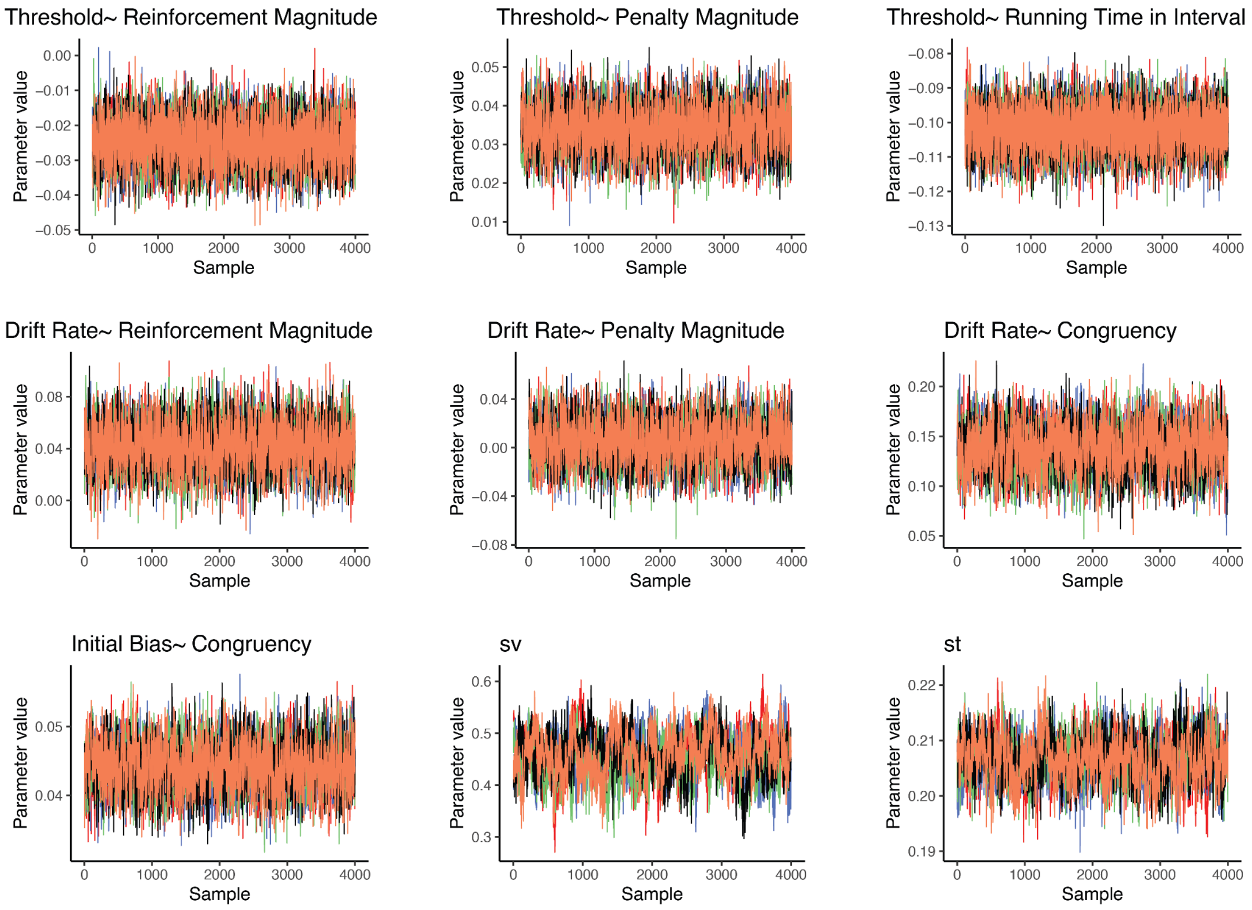


Figure S7 shows the trace plots for the Protector Game model only for each of the parameters. The Trace plots visualize the parameter value selection and look like lovely fuzzy caterpillars indicating good convergence.

**Figure S8.** Posterior Predictive Checks for Protector only Model

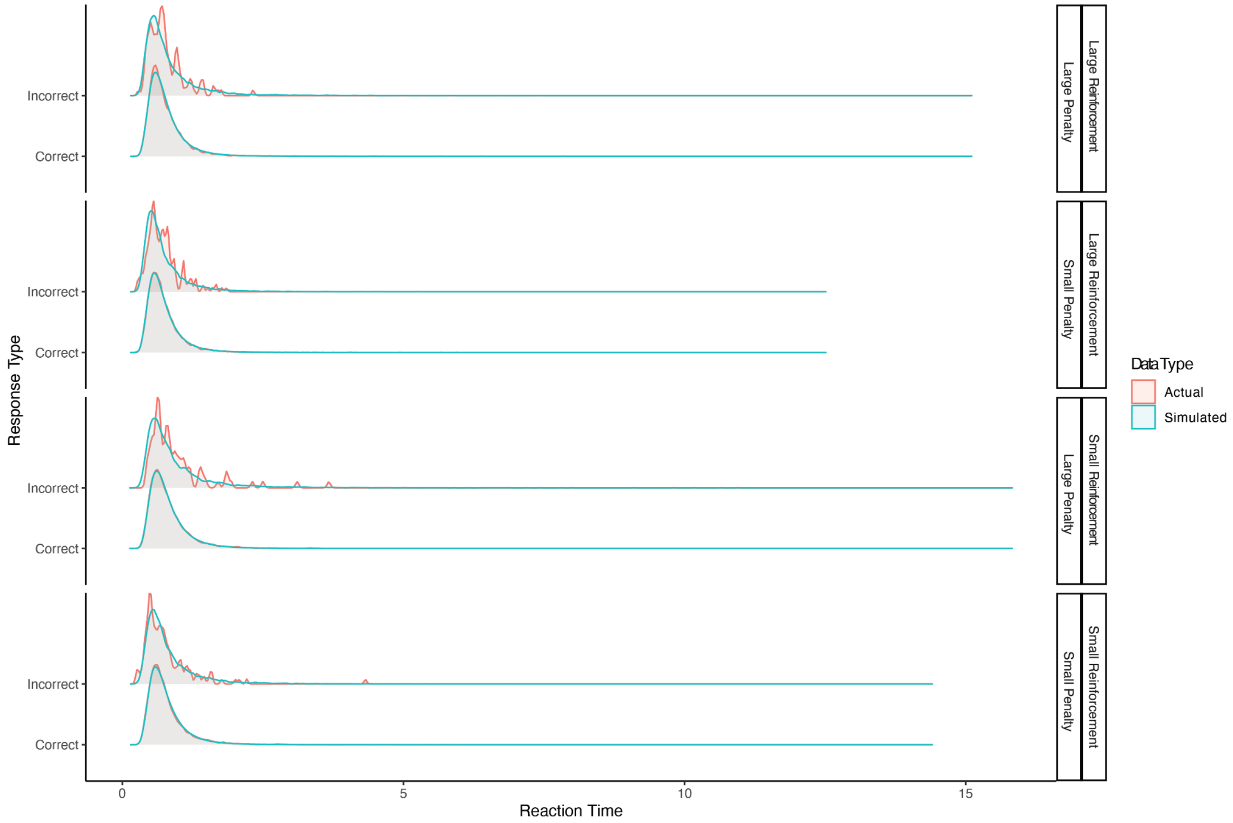


We performed a posterior predictive check for the fitted Drift Diffusion Model (DDM). We generated 500 independent samples from the posterior distribution of fitted parameters and then simulated the reaction time distribution for each posterior sample. The predicted reaction time distribution matches with the actual reaction time distribution for each condition.

**Figure S9.** Threshold and Drift Rates by Condition Estimated from Models fit on accuracy and reaction times from Collector and Protector Games Separately

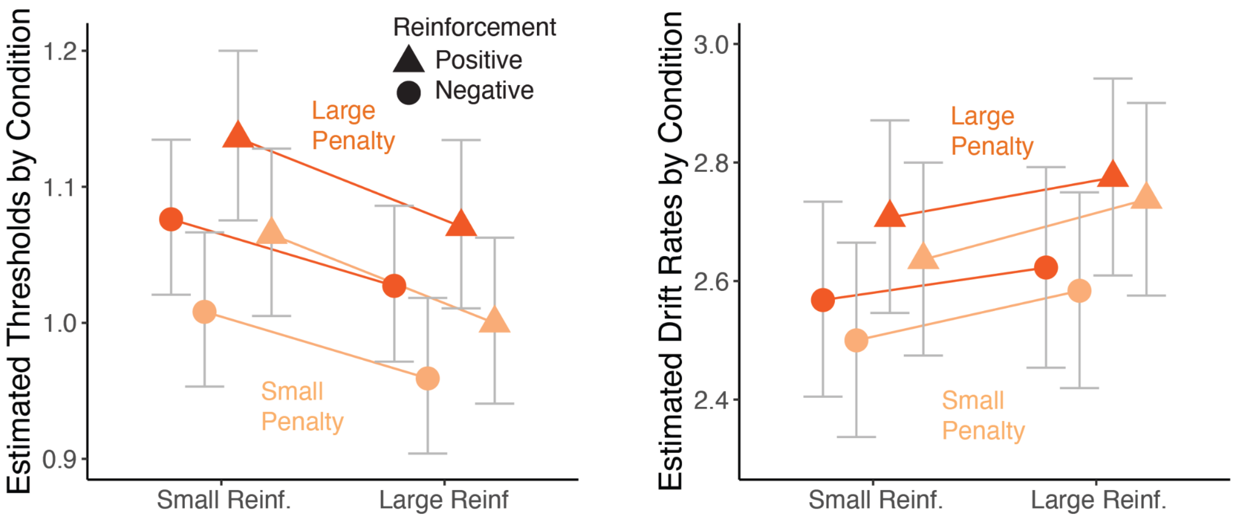

### Ratings

#### **Table S8.** Ratings within the Collector Game

Table S8 Shows the details of the fitted mixed models for the subjective ratings of Pleasantness, Arousal, Difficulty, Motivation, Effort, and Attention

| **Collector** | | | | | | | | | | | | |
|---|---|---|---|---|---|---|---|---|---|---|---|---|
| | **Pleasant** | | | | **Arousal** | | | | **Difficulty** | | | |
| *Predictors* | *Estimates* | *CI* | *Statistic* | *p* | *Estimates* | *CI* | *Statistic* | *p* | *Estimates* | *CI* | *Statistic* | *p* |
| Intercept | 0.53 | 0.34 – 0.72 | 5.39 | **<0.001** | 1.11 | 0.85 – 1.37 | 8.45 | **<0.001** | 0.39 | 0.12 – 0.67 | 2.84 | **0.005** |
| Reinforcement Magnitude | 1.71 | 1.52 – 1.91 | 17.53 | **<0.001** | 0.44 | 0.18 – 0.70 | 3.33 | **0.001** | -0.06 | -0.33 – 0.21 | -0.42 | 0.675 |
| Penalty Magnitude | -1.32 | -1.52 – -1.13 | -13.55 | **<0.001** | 0.62 | 0.36 – 0.88 | 4.71 | **<0.001** | 0.84 | 0.57 – 1.11 | 6.05 | **<0.001** |
| Age | -0.03 | -0.22 – 0.17 | -0.29 | 0.775 | 0.02 | -0.24 – 0.28 | 0.13 | 0.893 | -0.37 | -0.65 – -0.09 | -2.61 | **0.009** |
| Sex | -0.02 | -0.21 – 0.18 | -0.17 | 0.865 | 0.16 | -0.10 – 0.42 | 1.25 | 0.214 | 0.03 | -0.25 – 0.30 | 0.19 | 0.846 |
| Reinf. Magnitude x Penalty Magnitude | 0.05 | -0.14 – 0.24 | 0.49 | 0.623 | -0.26 | -0.52 – -0.00 | -1.97 | **0.050** | -0.20 | -0.47 – 0.07 | -1.45 | 0.149 |
| Observations | 364 | | | | 364 | | | | 364 | | | |
| $R^2$ / $R^2$ adjusted | 0.588 / 0.582 | | | | 0.104 / 0.091 | | | | 0.121 / 0.108 | | | |

| | **Motivation** | | | | **Effort** | | | | **Attention** | | | |
|---|---|---|---|---|---|---|---|---|---|---|---|---|
| *Predictors* | *Estimates* | *CI* | *Statistic* | *p* | *Estimates* | *CI* | *Statistic* | *p* | *Estimates* | *CI* | *Statistic* | *p* |
| Intercept | 2.32 | 2.09 – 2.56 | 19.41 | **<0.001** | 3.13 | 2.93 – 3.32 | 30.93 | **<0.001** | 2.10 | 1.80 – 2.40 | 13.85 | **<0.001** |
| Reinforcement Magnitude | 0.96 | 0.72 – 1.19 | 8.03 | **<0.001** | 0.51 | 0.31 – 0.71 | 5.02 | **<0.001** | 0.39 | 0.11 – 0.66 | 2.76 | **0.007** |
| Penalty Magnitude | 0.28 | 0.04 – 0.51 | 2.33 | **0.020** | 0.43 | 0.23 – 0.63 | 4.26 | **<0.001** | 0.31 | 0.03 – 0.59 | 2.19 | **0.030** |
| Age | 0.20 | -0.04 – 0.43 | 1.65 | 0.100 | 0.37 | 0.17 – 0.57 | 3.66 | **<0.001** | -0.04 | -0.44 – 0.37 | -0.18 | 0.859 |
| Sex | 0.04 | -0.20 – 0.27 | 0.30 | 0.761 | 0.03 | -0.17 – 0.23 | 0.27 | 0.789 | 0.00 | -0.28 – 0.29 | 0.02 | 0.982 |
| Reinf. Magnitude x Penalty Magnitude | -0.54 | -0.78 – -0.31 | -4.54 | **<0.001** | -0.36 | -0.55 – -0.16 | -3.53 | **<0.001** | -0.37 | -0.65 – -0.10 | -2.65 | **0.009** |
| Observations | 364 | | | | 364 | | | | 170 | | | |
| $R^2$ / $R^2$ adjusted | 0.211 / 0.200 | | | | 0.153 / 0.141 | | | | 0.125 / 0.098 | | | |

#### **Table S9.** Ratings within the Protector Game

Table S9 Shows the details of the fitted mixed models for the subjective ratings of Pleasantness, Arousal, Difficulty, Motivation, Effort, and Attention

| **Protector Ratings** | | | | | | | | | | | | |
|---|---|---|---|---|---|---|---|---|---|---|---|---|
| | **Pleasant** | | | | **Arousal** | | | | **Difficulty** | | | |
| *Predictors* | *Estimates* | *CI* | *Statistic* | *p* | *Estimates* | *CI* | *Statistic* | *p* | *Estimates* | *CI* | *Statistic* | *p* |
| Intercept | 0.08 | -0.12 – 0.29 | 0.80 | 0.422 | 1.10 | 0.85 – 1.35 | 8.69 | **<0.001** | 0.43 | 0.14 – 0.72 | 2.93 | **0.004** |
| Reinforcement Magnitude | 1.66 | 1.46 – 1.87 | 15.95 | **<0.001** | 0.38 | 0.13 – 0.62 | 2.98 | **0.003** | -0.05 | -0.34 – 0.23 | -0.36 | 0.716 |
| Penalty Magnitude | -1.34 | -1.54 – -1.13 | -12.83 | **<0.001** | 0.64 | 0.39 – 0.89 | 5.07 | **<0.001** | 0.67 | 0.38 – 0.96 | 4.58 | **<0.001** |
| Sex | -0.07 | -0.28 – 0.14 | -0.66 | 0.509 | 0.07 | -0.18 – 0.32 | 0.52 | 0.604 | 0.20 | -0.09 – 0.49 | 1.36 | 0.174 |
| Reinf. Magnitude x Penalty Magnitude | 0.06 | -0.15 – 0.26 | 0.55 | 0.580 | -0.23 | -0.47 – 0.02 | -1.80 | 0.073 | -0.23 | -0.52 – 0.06 | -1.56 | 0.120 |
| Observations | 364 | | | | 364 | | | | 364 | | | |
| $R^2$ / $R^2$ adjusted | 0.540 / 0.534 | | | | 0.101 / 0.089 | | | | 0.079 / 0.067 | | | |

| | **Motivation** | | | | **Effort** | | | | **Attention** | | | |
|---|---|---|---|---|---|---|---|---|---|---|---|---|
| *Predictors* | *Estimates* | *CI* | *Statistic* | *p* | *Estimates* | *CI* | *Statistic* | *p* | *Estimates* | *CI* | *Statistic* | *p* |
| Intercept | 1.84 | 1.57 – 2.11 | 13.42 | **<0.001** | 2.97 | 2.75 – 3.18 | 27.15 | **<0.001** | 5.00 | 5.00 – 5.00 | | |
| Reinforcement Magnitude | 1.01 | 0.75 – 1.28 | 7.41 | **<0.001** | 0.41 | 0.20 – 0.63 | 3.78 | **<0.001** | 0.00 | 0.00 – 0.00 | | |
| Penalty Magnitude | 0.30 | 0.03 – 0.57 | 2.22 | **0.027** | 0.43 | 0.21 – 0.64 | 3.91 | **<0.001** | 0.00 | 0.00 – 0.00 | | |
| Age | 0.28 | 0.01 – 0.55 | 2.03 | **0.043** | 0.31 | 0.10 – 0.52 | 2.86 | **0.004** | 0.00 | 0.00 – 0.00 | | |
| Sex | -0.16 | -0.43 – 0.11 | -1.19 | 0.236 | -0.05 | -0.26 – 0.17 | -0.45 | 0.656 | 0.00 | 0.00 – 0.00 | | |
| Reinf. Magnitude x Penalty Magnitude | -0.59 | -0.86 – -0.32 | -4.31 | **<0.001** | -0.35 | -0.57 – -0.14 | -3.23 | **0.001** | -0.00 | -0.00 – 0.00 | | |
| Observations | 364 | | | | 364 | | | | 340 | | | |
| $R^2$ / $R^2$ adjusted | 0.195 / 0.183 | | | | 0.116 / 0.104 | | | | NA | | | |

#### **Table S10.** Ratings within the Collector and Protector Game

Table S10 Shows the details of the fitted mixed models for the subjective ratings of Pleasantness, Arousal, Difficulty, Motivation, Effort, and Attention

| **Ratings Combined Across Games** | | | | | | | | | | | | |
|---|---|---|---|---|---|---|---|---|---|---|---|---|
| | **Pleasant** | | | | **Arousal** | | | | **Difficulty** | | | |
| *Predictors* | *Estimates* | *CI* | *Statistic* | *p* | *Estimates* | *CI* | *Statistic* | *p* | *Estimates* | *CI* | *Statistic* | *p* |
| Intercept | 0.31 | 0.17 – 0.45 | 4.36 | **<0.001** | 1.11 | 0.93 – 1.28 | 12.13 | **<0.001** | 0.41 | 0.22 – 0.61 | 4.10 | **<0.001** |
| Reinforcement Magnitude | 1.69 | 1.55 – 1.83 | 23.88 | **<0.001** | 0.41 | 0.23 – 0.59 | 4.48 | **<0.001** | -0.06 | -0.25 – 0.14 | -0.56 | 0.578 |
| Penalty Magnitude | -1.34 | -1.47 – -1.20 | -18.87 | **<0.001** | 0.63 | 0.45 – 0.81 | 6.90 | **<0.001** | 0.76 | 0.56 – 0.95 | 7.51 | **<0.001** |
| Reinforcement Valence | 0.22 | 0.08 – 0.36 | 3.12 | **0.002** | 0.01 | -0.16 – 0.19 | 0.15 | 0.880 | -0.02 | -0.22 – 0.17 | -0.24 | 0.807 |
| Age | 0.03 | -0.11 – 0.17 | 0.49 | 0.628 | -0.01 | -0.19 – 0.16 | -0.16 | 0.871 | -0.33 | -0.53 – -0.12 | -3.16 | **0.002** |
| Sex | -0.04 | -0.18 – 0.10 | -0.56 | 0.574 | 0.11 | -0.06 – 0.29 | 1.25 | 0.210 | 0.11 | -0.09 – 0.31 | 1.10 | 0.270 |
| Reinf. Magnitude x Penalty Magnitude | 0.05 | -0.09 – 0.19 | 0.75 | 0.453 | -0.24 | -0.42 – -0.06 | -2.67 | **0.008** | -0.21 | -0.41 – -0.02 | -2.13 | **0.033** |
| Reinf. Magnitude x Reinf. Valence | 0.02 | -0.12 – 0.16 | 0.23 | 0.818 | 0.03 | -0.15 – 0.21 | 0.35 | 0.730 | -0.00 | -0.20 – 0.19 | -0.03 | 0.977 |
| Penalty Magnitude x Reinf. Valence | 0.02 | -0.12 – 0.16 | 0.28 | 0.779 | -0.01 | -0.19 – 0.17 | -0.14 | 0.889 | 0.08 | -0.12 – 0.28 | 0.81 | 0.417 |
| Reinf. Magnitude x Penalty Magnitude x Reinf. Valence | -0.00 | -0.14 – 0.14 | -0.05 | 0.964 | -0.02 | -0.20 – 0.16 | -0.21 | 0.837 | 0.01 | -0.18 – 0.21 | 0.14 | 0.886 |
| Observations | 728 | | | | 728 | | | | 728 | | | |
| $R^2$ / $R^2$ adjusted | 0.570 / 0.565 | | | | 0.102 / 0.091 | | | | 0.099 / 0.087 | | | |

| | **Motivation** | | | | **Effort** | | | | **Attention** | | | |
|---|---|---|---|---|---|---|---|---|---|---|---|---|
| *Predictors* | *Estimates* | *CI* | *Statistic* | *p* | *Estimates* | *CI* | *Statistic* | *p* | *Estimates* | *CI* | *Statistic* | *p* |
| Intercept | 2.08 | 1.90 – 2.26 | 22.86 | **<0.001** | 3.04 | 2.90 – 3.19 | 40.93 | **<0.001** | 3.68 | 3.55 – 3.81 | 55.48 | **<0.001** |
| Reinforcement Magnitude | 0.99 | 0.81 – 1.17 | 10.92 | **<0.001** | 0.46 | 0.31 – 0.61 | 6.20 | **<0.001** | 0.31 | 0.18 – 0.44 | 4.72 | **<0.001** |
| Penalty Magnitude | 0.29 | 0.11 – 0.47 | 3.21 | **0.001** | 0.43 | 0.28 – 0.57 | 5.77 | **<0.001** | 0.32 | 0.19 – 0.45 | 4.80 | **<0.001** |
| Reinforcement Valence | 0.21 | 0.04 – 0.39 | 2.35 | **0.019** | 0.07 | -0.08 – 0.22 | 0.93 | 0.352 | 0.04 | -0.09 – 0.17 | 0.55 | 0.583 |
| Age | 0.24 | 0.06 – 0.42 | 2.63 | **0.009** | 0.34 | 0.19 – 0.49 | 4.59 | **<0.001** | 0.28 | 0.15 – 0.41 | 4.24 | **<0.001** |
| Sex | -0.06 | -0.24 – 0.11 | -0.71 | 0.479 | -0.01 | -0.16 – 0.14 | -0.15 | 0.882 | -0.29 | -0.42 – -0.16 | -4.38 | **<0.001** |

| | | | | | | | | | | | | |
|---|---|---|---|---|---|---|---|---|---|---|---|---|
| Reinf. Magnitude x Penalty Magnitude | -0.57 | -0.75 – -0.39 | -6.26 | **<0.001** | -0.35 | -0.50 – -0.21 | -4.77 | **<0.001** | -0.28 | -0.41 – -0.15 | -4.26 | **<0.001** |
| Reinf. Magnitude x Reinf. Valence | -0.02 | -0.20 – 0.16 | -0.22 | 0.827 | 0.05 | -0.10 – 0.19 | 0.64 | 0.520 | 0.07 | -0.06 – 0.20 | 1.00 | 0.319 |
| Penalty Magnitude x Reinf. Valence | -0.03 | -0.20 – 0.15 | -0.28 | 0.779 | -0.00 | -0.15 – 0.15 | -0.01 | 0.993 | 0.03 | -0.10 – 0.16 | 0.41 | 0.682 |
| Reinf. Magnitude x Penalty Magnitude x Reinf. Valence | 0.04 | -0.14 – 0.22 | 0.41 | 0.682 | -0.00 | -0.15 – 0.15 | -0.00 | 0.999 | -0.06 | -0.19 – 0.07 | -0.91 | 0.362 |
| Observations | 728 | | | | 728 | | | | 728 | | | |
| $R^2$ / $R^2$ adjusted | 0.207 / 0.197 | | | | 0.135 / 0.124 | | | | 0.111 / 0.100 | | | |

#### **Table S11** Robust regression model results for the composite motivation ratings for the Collector Game

| **Collector** | | | | |
|---|---|---|---|---|
| | **Composite Motivation Score** | | | |
| *Predictors* | *Estimates* | *CI* | *Statistic* | *p* |
| Intercept | 2.98 | 2.81 – 3.15 | 34.45 | **<0.001** |
| Reinforcement Magnitude | 0.60 | 0.43 – 0.77 | 6.97 | **<0.001** |
| Penalty Magnitude | 0.37 | 0.20 – 0.54 | 4.23 | **<0.001** |
| Age | 0.33 | 0.16 – 0.50 | 3.79 | **<0.001** |
| Sex | -0.11 | -0.28 – 0.06 | -1.27 | 0.205 |
| Reinf. Magnitude x Penalty Magnitude | -0.39 | -0.56 – -0.22 | -4.57 | **<0.001** |
| Observations | 364 | | | |
| $R^2$ / $R^2$ adjusted | 0.218 / 0.207 | | | |

#### **Table S12** Robust regression model results for the composite motivation ratings for the Protector Game

| **Protector Game** | | | | |
|---|---|---|---|---|
| | **Composite Motivation Score** | | | |
| *Predictors* | *Estimates* | *CI* | *Statistic* | *p* |

| | | | | |
|---|---|---|---|---|
| Intercept | 2.76 | 2.58 – 2.94 | 29.59 | **<0.001** |
| Reinforcement Magnitude | 0.57 | 0.38 – 0.75 | 6.07 | **<0.001** |
| Penalty Magnitude | 0.36 | 0.17 – 0.54 | 3.83 | **<0.001** |
| Age | 0.28 | 0.10 – 0.47 | 3.03 | **0.003** |
| Sex | -0.17 | -0.36 – 0.01 | -1.86 | 0.064 |
| Reinf. Magnitude x Penalty Magnitude | -0.37 | -0.56 – -0.19 | -4.02 | **<0.001** |
| Observations | 364 | | | |
| $R^2$ / $R^2$ adjusted | 0.181 / 0.170 | | | |

#### **Table S13** Robust regression model results for the composite motivation ratings for the Collector and Protector Game

| **All Ratings** | | | | |
|---|---|---|---|---|
| | **Composite Motivation Score** | | | |
| *Predictors* | *Estimates* | *CI* | *Statistic* | *p* |
| Intercept | 2.87 | 2.75 – 3.00 | 45.20 | **<0.001** |
| Reinforcement Magnitude | 0.58 | 0.46 – 0.71 | 9.21 | **<0.001** |
| Penalty Magnitude | 0.36 | 0.24 – 0.49 | 5.69 | **<0.001** |
| Reinforcement Valence | 0.10 | -0.02 – 0.22 | 1.58 | 0.114 |
| Age | 0.31 | 0.18 – 0.43 | 4.80 | **<0.001** |
| Sex | -0.14 | -0.27 – -0.02 | -2.23 | **0.026** |
| Reinf. Magnitude x Penalty Magnitude | -0.38 | -0.51 – -0.26 | -6.06 | **<0.001** |
| Reinf. Magnitude x Reinf. Valence | 0.02 | -0.11 – 0.14 | 0.29 | 0.774 |
| Penalty Magnitude x Reinf. Valence | 0.00 | -0.12 – 0.13 | 0.02 | 0.981 |
| Reinf. Magnitude x Penalty Magnitude x Reinf. Valence | -0.01 | -0.13 – 0.12 | -0.14 | 0.887 |
| Observations | 728 | | | |
| $R^2$ / $R^2$ adjusted | 0.201 / 0.191 | | | |

**Figure S10**. Correlations of Ratings for Cues

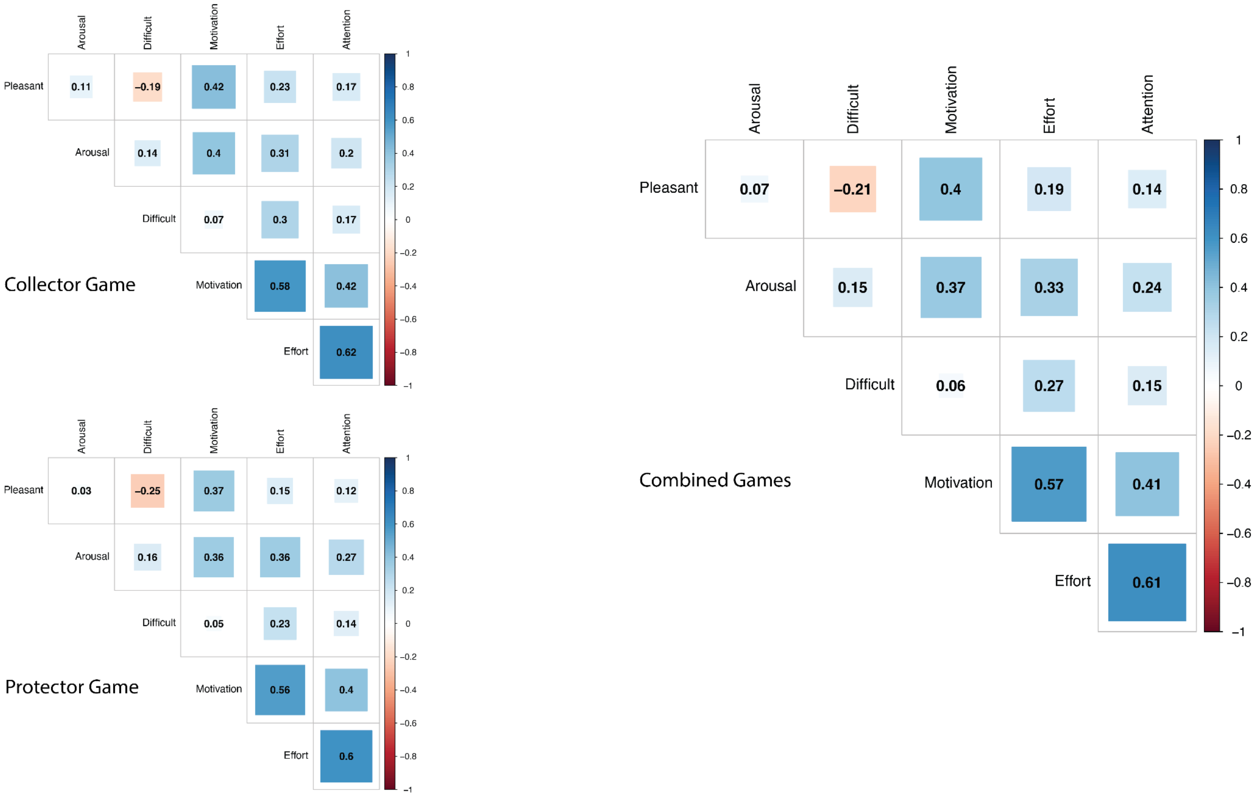


Top Left Collector Game: Positive correlation between Motivation and Effort of 0.58 (t(362) = 13.635, p <0.001). Positive correlation between Motivation and Attention of 0.42 (t(362) = 8.735, p <0.001). Positive correlation between Attention and Effort of 0.62 (t(362) = 15.011, p <0.001 )
Bottom Left Collector Game: Positive correlation between Motivation and Effort of 0.56 (t(362) = 12.757, p <0.001). Positive correlation between Motivation and Attention of 0.40 (t(362) = 8.336, p <0.001). Positive correlation between Attention and Effort of 0.60 (t(362) = 14.403, p <.001 ).
Right Combined Games: Positive correlation between Motivation and Effort of 0.57 (t(726) = 18.584, p <0.001). Positive correlation between Motivation and Attention of 0.41 (t(726) = 12.075, p <0.001). Positive correlation between Attention and Effort of 0.61 (t(726) = 20.819, p <.001 )
